# Optimal Evidence Accumulation On Social Networks

**DOI:** 10.1101/442848

**Authors:** Bhargav Karamched, Simon Stolarczyk, Zachary P. Kilpatrick, Krešimir Jossić

**Affiliations:** Department of Mathematics, University of Houston, Houston TX 77204; Department of Applied Mathematics, University of Colorado, Boulder CO 80309; Departments of Mathematics, and Biology and Biochemistry, University of Houston, Houston TX 77204, Department of BioSciences, Rice University, Houston TX 77005

**Keywords:** decision-making, probabilistic inference, social networks

## Abstract

A fundamental question in biology is how organisms integrate sensory and social evidence to make decisions. However, few models describe how both these streams of information can be combined to optimize choices. Here we develop a normative model for collective decision making in a network of agents performing a two-alternative forced choice task. We assume that rational (Bayesian) agents in this network make private measurements, and observe the decisions of their neighbors until they accumulate sufficient evidence to make an irreversible choice. As each agent communicates its decision to those observing it, the flow of social information is described by a directed graph. The decision-making process in this setting is intuitive, but can be complex. We describe when and how the *absence* of a decision of a neighboring agent communicates social information, and how an agent must marginalize over all unobserved decisions. We also show how decision thresholds and network connectivity affect group evidence accumulation, and describe the dynamics of decision making in social cliques. Our model provides a bridge between the abstractions used in the economics literature and the evidence accumulator models used widely in neuroscience and psychology.

## 1. Introduction

Understanding how organisms use sensory and social information to make decisions is of fundamental interest in biology, sociology, and economics [13, 16, 20]. Psychologists and neuroscientists have developed a variety of experimental approaches to probe how humans and other animals make choices. Particularly popular are variants of the two-alternative forced choice task (2AFC) where an observer is asked to decide between two options based on information obtained from one or more noisy observations [7, 24, 47, 48]. The 2AFC task has motivated several mathematical models that successfully explain how humans use sequentially-presented evidence to make decisions [9, 15, 55].

Most evidence accumulation models take the form of drift-diffusion stochastic processes that describe the information gathered by lone observers [7]. However, humans and many other animals do not live in isolation and make decisions based on more than their own private observations. Animals watch each other as they forage [29, 34]. Stock traders, while not privy to all of their competitor’s information, can still observe each other’s decisions. To make the best choices many biological agents thus also take into account the observed choices of others [37, 42, 45].

Here we address the question of how idealized agents in a social network should combine a sequence of private measurements with observed decisions of other individuals to choose between two options. We refer to the information an agent receives from its neighbors as *social information*, and information available only to the agent as *private information*. As agents do not share their private information with others directly, they only reveal their beliefs through their choices. These choices are based on private and social information, and thus reveal something about the total evidence an agent has collected.

We assume that private measurements, and, as a consequence, observed choices can improve the odds of making the right decision. However neither type of information affords certainty about which choice is correct. We take a probabilistic (Bayesian) approach to describe the behavior of rational agents who make *immutable* decisions once they have accrued sufficient evidence. Their choices thus provide information about their beliefs at a single point in time.

There are two reasons for assuming that agents only share their decision: First, many organisms communicate their decisions, but not the evidence they used to reach them. For example, herding animals in motion can only communicate their chosen direction of movement [14,40]. Animals that forage in groups may communicate their preferred feeding locations [19, 49], but not the evidence, or the process which they used to decide. Human traders can see their competitor’s choice to buy or sell a stock, but may not have access to the evidence that lead to these actions. Second, if agents communicate all information they gathered to their neighbors, the problem is mathematically trivial as every agent obtains all evidence provided to the entire network.

The behavior and performance of rational agents in a social network can depend sensitively on how information is shared [1, 35]. Surprisingly, in some cases rational agents perform better in isolation, on average. This can happen when social information from a small subset of agents dominates the decisions of the collective, as in classical examples of *herding behavior* [5, 21]. On the other hand, if after a single observation agents share their belief and continue to do so repeatedly in a recurrent network, then all agents can asymptotically come to agree on the most probable choice given the totality of evidence obtained by the network [2, 22, 23, 35]. In this case of *asymptotic learning*, all agents are eventually able to make use of all the private information in the network by observing the preferences of their neighbors.

In contrast the decisions of agents in our model are immutable. As a consequence, asymptotic learning typically does not occur. We also show how a rational agent who only observes a portion of the network needs to marginalize over the decisions of all unobserved agents to correctly integrate an observed decision. Interestingly when decision thresholds are asymmetric, even the absence of a decision can communicate information deterministically until a decision is reached. In recurrent networks observing the absence of a decision by a neighbor who has observed your own indecision can lead to additional social information, and this can happen repeatedly. This is akin to situations addressed in the literature on *common knowledge* [2, 22, 33].

Our evidence accumulation model shows how to best combine streams of private, and social evidence to make decisions. Such evidence accumulation models have been shown to provide excellent description of decision making in various tasks, and there is evidence that they can explain the formation of social decisions [29]. They may thus point to common mechanisms for human decision-making.

## 2. Definitions and setup

We consider a set of agents who accumulate evidence to decide between two states, *H*^+^ or *H^−^*. Each agent is rational (Bayesian): They compute and compare the probability of each state (or hypothesis), based on all evidence they accrue. We assume that agents make a decision once the conditional probability of one of the states, given all the accumulated observations, crosses a predetermined threshold [7, 59].

To make a choice, agents gather both private (Priv) and social (Soc) information. Private information comes from a sequence of noisy observations (measurements) of the state, *H* ∈ {*H*^+^*, H^−^*}. Agents also gather social information by observing each other’s decisions, as in previous models of foraging animal groups [34], consumer networks [1], and opinion exchange on social networks [60]. Thus agents do not share their private measurements directly, but only observe whether their neighbors have made a choice, and, if so, what that choice was.

### Evidence accumulation for a single agent

The problem of a single agent accumulating private evidence to decide between two options has been thoroughly studied [7, 24, 46, 47, 55, 56, 59]. Assume that an agent makes a sequence of noisy observations, *ξ*_1:*t*_ with *ξ_i_* ∈ Ξ*, i* ∈ {1*, …, t*}, and Ξ ⊂ ℝ finite. The observations *ξ_i_* are independent and identically distributed, conditioned on the true state *H* ∈ {*H*^+^*, H^−^*},

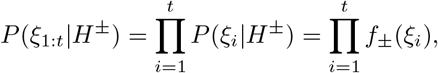

where *f_±_*(*ξ*) := *P* (*ξ|H^±^*) are the measurement distributions, which are probability mass functions since Ξ is finite. Observations *ξ* are drawn from the same set Ξ in either state *H^±^*, and states are distinguished by the different measurement distributions *f_±_*(*ξ*).

To compute *P* (*H|ξ*_1:*t*_) the agent uses Bayes’ Rule: For simplicity, we assume that the agent knows the measurement distributions *f_±_*(*ξ*), uses a flat prior, *P* (*H*^+^) = *P* (*H^−^*) = 1*/*2, and that observations are conditionally independent. Thus, the log likelihood ratio (LLR) of the two states at time *t* is

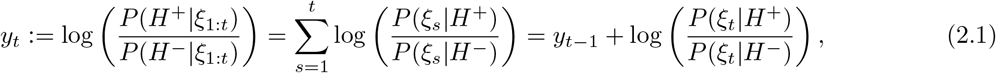

where *y*_0_ = 0, since both states are equally likely *a priori*, log(*P* (*H*^+^)*/P* (*H^−^*)) = 0. We also refer to the LLR as the *belief* of the agent.

An ideal agent continues making observations while *θ_−_ < y_t_ < θ*_+_, and makes a decision after acquiring sufficient evidence, choosing *H*^+^ (*H^−^*) once *y_t_ ≥ θ*_+_ (*y_t_ ≤ θ_−_*). We assume *θ_−_ <* 0 *< θ*_+_.

**Table 2.1.**
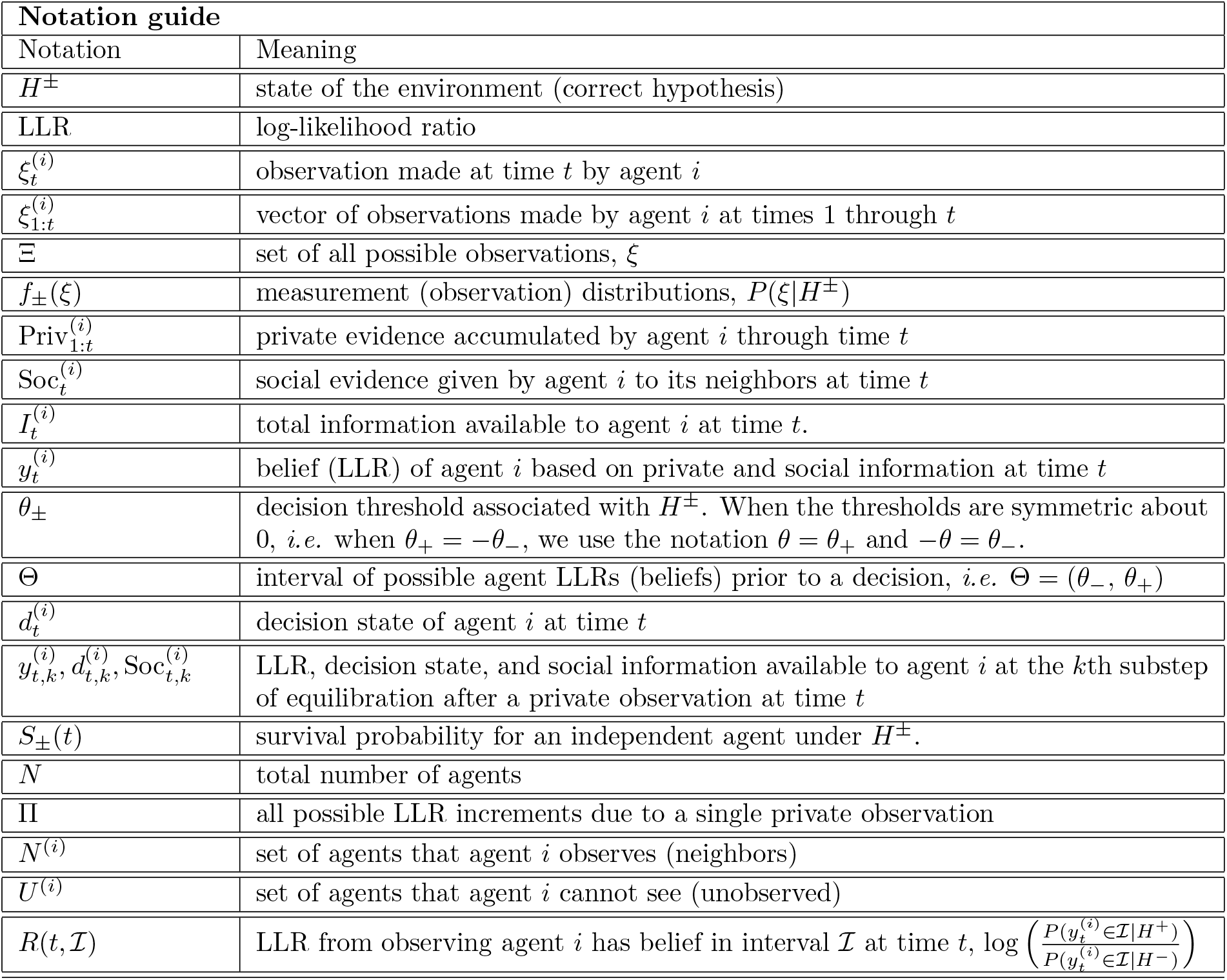
Notation guide.

## 3. Multiple agents

Limited bandwidth or physical distances between agents can limit their interactions [30, 41, 61]. We identify agents with a set of vertices, *V* = {1*, …, N*}, and communication between agents with a set of directed edges, *E,* between these vertices [25]. The agent at the tail of each directed edge communicates information to the agent at the edge’s head.

As in the case of a single observer, we assume that each agent, *i*, makes a sequence of noisy, identically-distributed measurements, 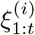, of the state, *H,* from a state–dependent distribution, 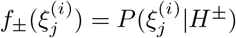. We assume that the observations are independent in time and between agents, conditioned on the state, 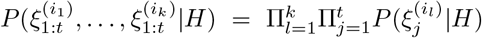 for any sequence of measurements, 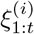, and set of agents, *i*_1_*, …, i_k_*, in the network. This conditional independence of the measurements is a strong assumption, and is unlikely to hold in practice. We return to this point in the Discussion.

An agent gathers social evidence by observing whether its neighbors have made a decision, and, once they have, what that decision is. Each agent thus gathers private and social evidence, and makes a decision when its belief (LLR) about the two states crosses one of the thresholds, *θ_−_ <* 0 *< θ*_+_. Importantly, *once an agent has made a decision, it cannot change it.* The absence of a decision thus communicates that an agent has not gathered sufficient evidence to make a choice, and hence that this agent’s belief (LLR) is still in the interval (*θ_−_, θ*_+_).

For simplicity, we assume that the distributions and thresholds, *θ_±_,* are identical across agents. The theory is similar if agents have different, but known, measurement distributions. The assumption that the measurement distributions, *f_±_*(*ξ*), are discrete, simplifies some convergence arguments. In subsequent work, we will extend our analysis to the case of continuous measurement distributions and time.

### Evidence accumulation with two agents

To illustrate how an agent integrates private and social information to reach a decision, we use the example network shown in Fig. 3.1a. We will show that even the absence of a decision can provide social information.

**Fig. 3.1.**
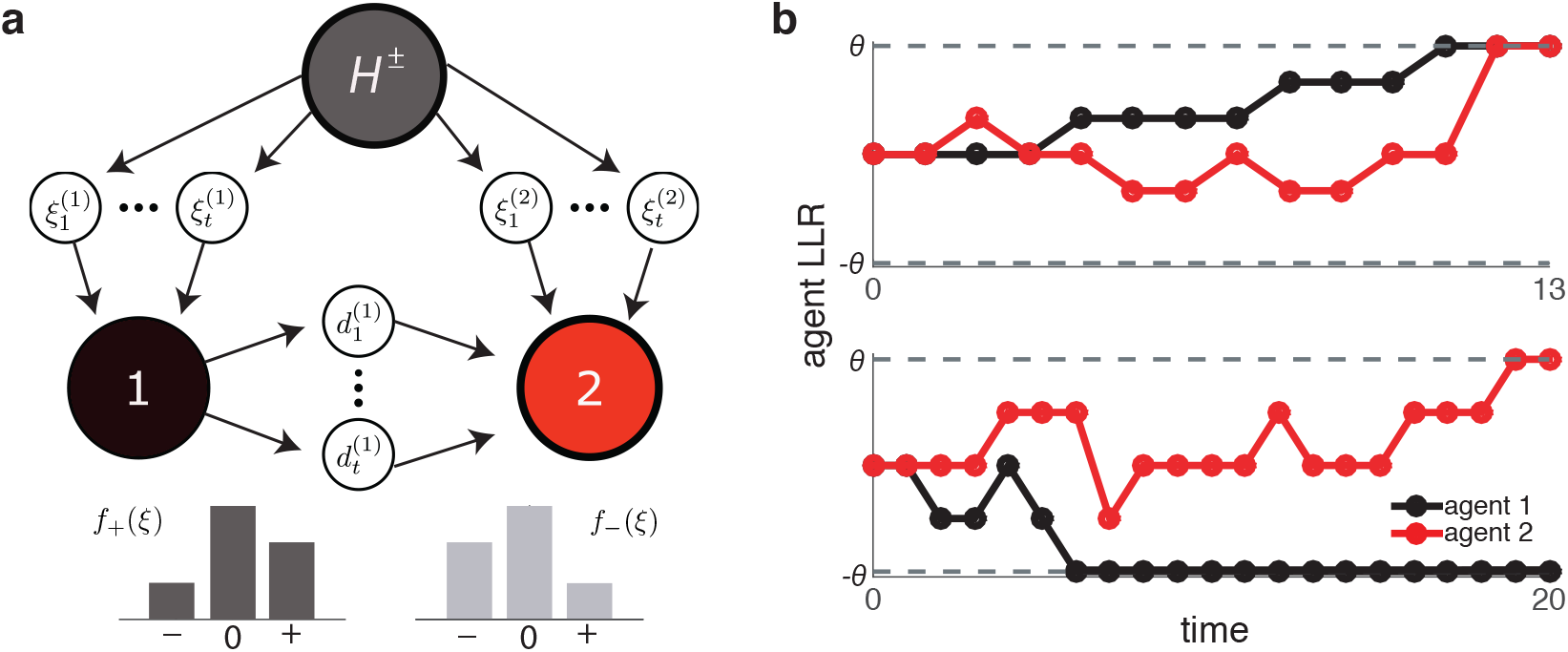
A pair of unidirectionally coupled agents deciding between the states H^+^ and H^−^ on the basis of private and social evidence. (a) Schematic of the information flow in the network. Agent 1 accumulates their own observations, 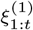, which result in a sequence of decision states, 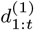, that is observed by agent 2. In addition, agent 2 gathers its own observations, 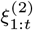, to make a decision. (b) Sample trajectories for the beliefs (LLRs) of the agents. Decisions are made when an agent’s belief crosses a threshold, θ_±_ = ±θ in this case. A decision of agent 1 leads to a jump in the belief of agent 2.

Let 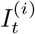 be the total information available to agent *i* at time *t*. The information available to agent 1, 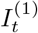, consists only of private observations. However, agent 2 makes private observations, and obtains social information from agent 1, both of which constitute 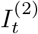. Agents base their decisions on the computed LLR, or belief, 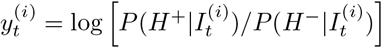. Each agent *i* makes a choice at the time *T*^(*i*)^ when 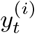 crosses one of the thresholds, *θ*_+_ or *θ_−_*.

Since 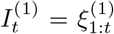, the belief of agent 1 is described by Eq. (2.1). At each time *t*, agent 2 observes the resulting *decision state*, 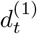, of agent 1 (Fig. 3.1a), where

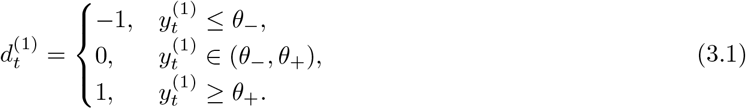

The decision state captures whether agent 1 made a decision by time *t*, and, if so, what that decision was.

Agent 2 also makes private observations, 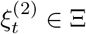. Assuming that the two agents make private observations synchronously, the information available to agent 2 at time *t* is 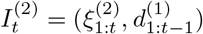: Decision 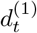 is thus not known to agent 2 until time *t* + 1. The observed decision state of agent 1 and the private observation, 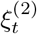, determine the belief of agent 2 at time *t*,

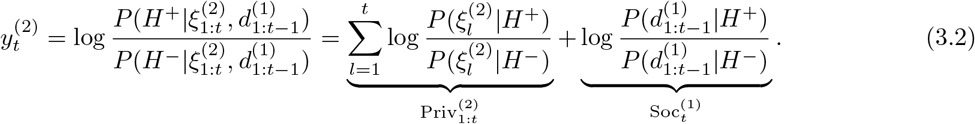

The last equality follows from our assumption that the observations are conditionally independent (See Proposition A.1 in Appendix A). The belief of agent 2 is thus a sum of the LLR corresponding to private observations 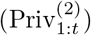, and the LLR corresponding to social information 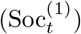. The stochastic process 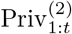 is defined in the same way for both agents, and is equivalent to that of a lone observer given in Eq. (2.1).

## 4. Social Information

We next ask how much evidence is provided by social information, that is, by observing the decision state of other agents in the directed network in Fig. 3.1a. We show that even observations of non-decisions, 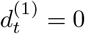, can be informative.

### Decision Evidence

As decisions are immutable, once an agent makes a choice, the social information it provides to its neighbors is fixed. Assume agent 1 chooses *H*^+^ at time *T*^(1)^. Then the belief of agent 2 at time *t > T*^(1)^ is:

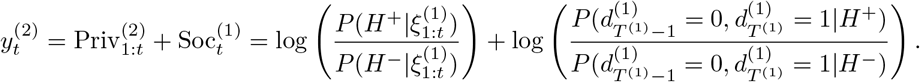

Intuitively, if, for instance, 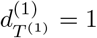, then agent 2 knows that 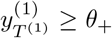. Agent 1 has reached its decision based on private observations, and hence none of the social information obtained by agent 2 is redundant. The belief of agent 1 at the time of the decision, *T*^(1)^ could exceed threshold. The evidence it has obtained may thus exceed *θ*_+_. However, this excess is small if the evidence obtained from each measurement is small.

The following proposition claims that, after observing a choice, agent 2 updates its belief by estimating the amount of evidence gathered by agent 1, *i.e.* by an amount close to *θ*_+_ (Fig. 3.1b).

#### Proposition 4.1.

*If* 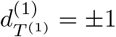, *then*

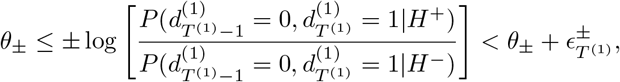

where

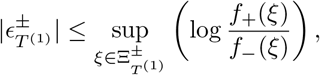

*and* 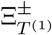 *is the set of all observations ξ_T_*_(1)_ that *trigger a H^±^ decision exactly at time T*^(1)^.

We prove Proposition 4.1 in Appendix B

### Social information prior to a decision

To understand why the absence of a decision can provide information, consider the case when *θ*_+_ ≠ *−θ_−_*. As one of the thresholds is closer to naught, in general each agent is more likely to choose one option over another by some time *t*. The absence of the more likely choice therefore reveals to an observer that the agent has gathered evidence favoring the alternative.

We will show that the lack of a decision is informative only in such asymmetric cases below.

#### DEFINITION 4.2.

*The measurement distributions P* (*ξ|H*^+^) = *f*_+_(*ξ*) *and P* (*ξ|H^−^*) = *f_−_*(*ξ*) *are* symmetric *if f*_+_(*−ξ*) = *f_−_*(*ξ*) *for every ξ* ∈ Ξ*. When θ*_+_ = *−θ_−_ we say that the* thresholds are symmetric.

It is frequently assumed, and experiments are frequently designed, so that threshold and measurement distributions are symmetric [50]. However, there are a number of interesting consequences when introducing asymmetries into the reward or measurement distributions [4], which suggest subjects adapt their priors or decision thresholds [27]. In examples we will assume that agents use asymmetric decisions thresholds (*θ*_+_ ≠ –*θ_−_*) due to a known asymmetry in the 2AFC task.

We call the social information agent 2 gathers before observing a choice by agent 1, the *non-decision evidence*. Using the decomposition in Eq. (3.2), we have

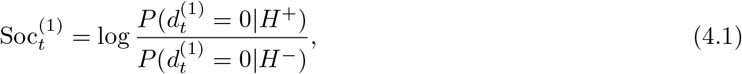

before agent 1 makes a decision. Let Θ := (*θ_−_, θ*_+_), and define the *survival probabilities* that the stochastically evolving belief remains within Θ as

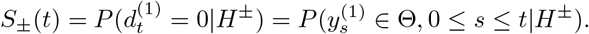

Then the social information provided by the absence of the decision by time *t* is,

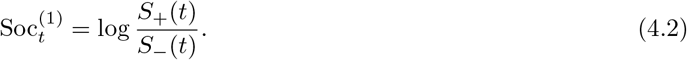

Note that if, for example, *S_−_*(*t*) *≤ S*_+_(*t*) for all *t ≥* 0, then *H^−^* decisions more often occur sooner than *H*^+^ decisions, and log[*S*_+_(*t*)*/S_−_*(*t*)] ≥ 0. Thus, observing that an agent has not made a choice by time *t* provides evidence in favor of the choice that requires more time to make.

Note also that social information depends only on the thresholds and measurement distributions of both agents, and not on the specific measurements of agent 2. Until agent 1 makes a decision, *the social information is deterministic*, *i.e.* independent of realization. We will see that a related result holds in more complex networks.

The next proposition shows that the symmetry of the thresholds and evidence distributions implies the absence of social information until a decision is made.

#### Proposition 4.3.

*If the measurement distributions and thresholds are symmetric then* 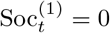 *when* 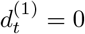.

*Proof*. Let *S*^(1)^(*t*) be the set of all observation sequences of agent 1, 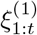, not leading to a decision. Then for all 0 *≤ s ≤ t*,

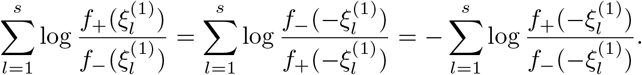

Here we have used the symmetry of measurement distributions in the first equality. Since the thresholds, and hence the interval Θ, are symmetric about 0, it follows that 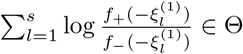, if and only if 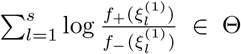. Therefore, 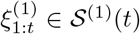, if and only if 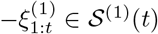. Moreover,

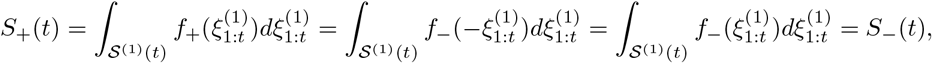

and thus observing the absence of a decision of agent 1 is uninformative, as

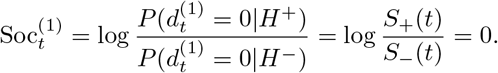

We show below that when measurement distributions and thresholds are symmetric, non-decisions are uninformative in any social network.

When thresholds and measurement distributions are not symmetric then typically *S_−_*(*t*) ≠ *S*_+_(*t*) and log [*S*_+_(*t*)*/S_−_*(*t*)] ≠ 0. Thus, the absence of a decision of agent 1 can provide evidence for one of the choices. We provide a concrete example next.

### Example: Belief as a Biased Random Walk

In the following example we chose the measurement distributions, *f*_±_(*ξ*), so that the belief increments due to private measurements satisfy log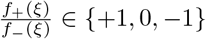 with respective probabilities {*p, s, q*}. This ensures that a decision based solely on private evidence results in a belief (LLR) exactly at threshold^1^. Beliefs then evolve as biased random walks on the integer lattice. Details of the model and the analysis are in Appendix C.

Agent 2 makes private observations and observes the decisions of agent 1, which are made only using private information. Agent 1 and 2’s beliefs are described by Eq. (2.1) and Eq. (3.2), respectively. Realizations of these processes are shown in Fig. 4.1. The social evidence obtained by agent 2 is independent of the particular realization, until the decision of agent 1, whereas private information is realization-dependent. When thresholds are small, an expression for social evidence can be obtained explicitly (See Appendix C).

**Fig. 4.1.**
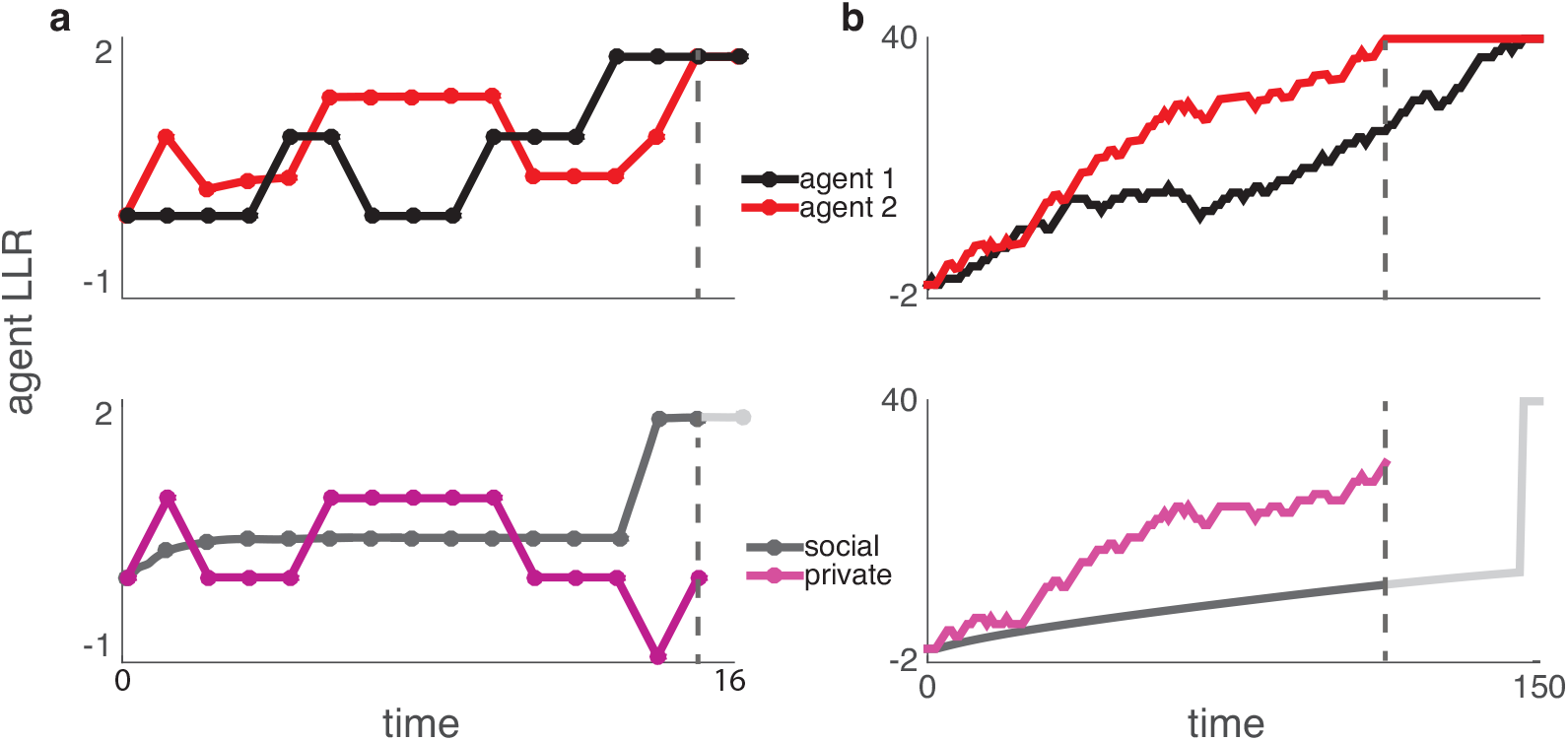
An example of the belief evolution in a two-agent, unidirectional network shown in Fig. 3.1. (a) The belief of agent 1 is a random walk on the integers. For boundaries at θ_−_ = −1 and θ_+_ = 2, initially two observations favoring H^+^ are required for decision 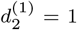. The belief of agent 2 is the sum of stochastically evolving private evidence, and deterministic social evidence until observing a choice. If agent 2 observes agent 1 choosing H^±^, its belief jumps by θ_±_. (b) The same processes with θ_−_ = –2 and θ_+_ = 40. The parameters used, and an analysis of the belief dynamics is given in Appendix C. Code to produce all the figures in the manuscript is available at https://github.com/Bargo727/NetworkDecisions.git.

First-step analysis shows that social evidence is not accrued significantly beyond the first few observations when decision thresholds are small. We show this for *θ_−_* = *−*1 and *θ*_+_ = 2 in Fig. 4.1a, with 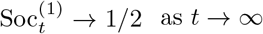 for our choice of parameter values (See Appendix C).

In general, social information, 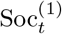, will converge in the direction of the larger threshold, in the absence of a decision. Intuitively, when |*θ*_+_|*>* |*θ_−_*| then more observations and a longer time are required for an agent’s belief to reach *θ*_+_, on average. In this case, observing that agent 1 has not chosen *H^−^* after a small initial period suggests that this agent has evidence favoring *H*^+^.

To illustrate the impact of asymmetry in the measurement distributions, we varied the probability, *p,* of an observation favoring *H*^+^, while keeping the increments in the belief fixed. When agent 2 observes the decisions of agent 1, the probability that both reach the correct decision is larger than when they both gather information independently (See Fig. 4.2a). In particular, as *p/q* decreases so that private observations provide less information, the impact of social information on accuracy is strengthened. With *θ_−_* small, little evidence is needed for an agent to choose *H^−^*. However, as *θ*_+_ is increased, social information pulls the belief of agent 2 in the direction of *θ*_+_ over time, but more evidence is required to reach the upper threshold *θ*_+_.

**Fig. 4.2.**
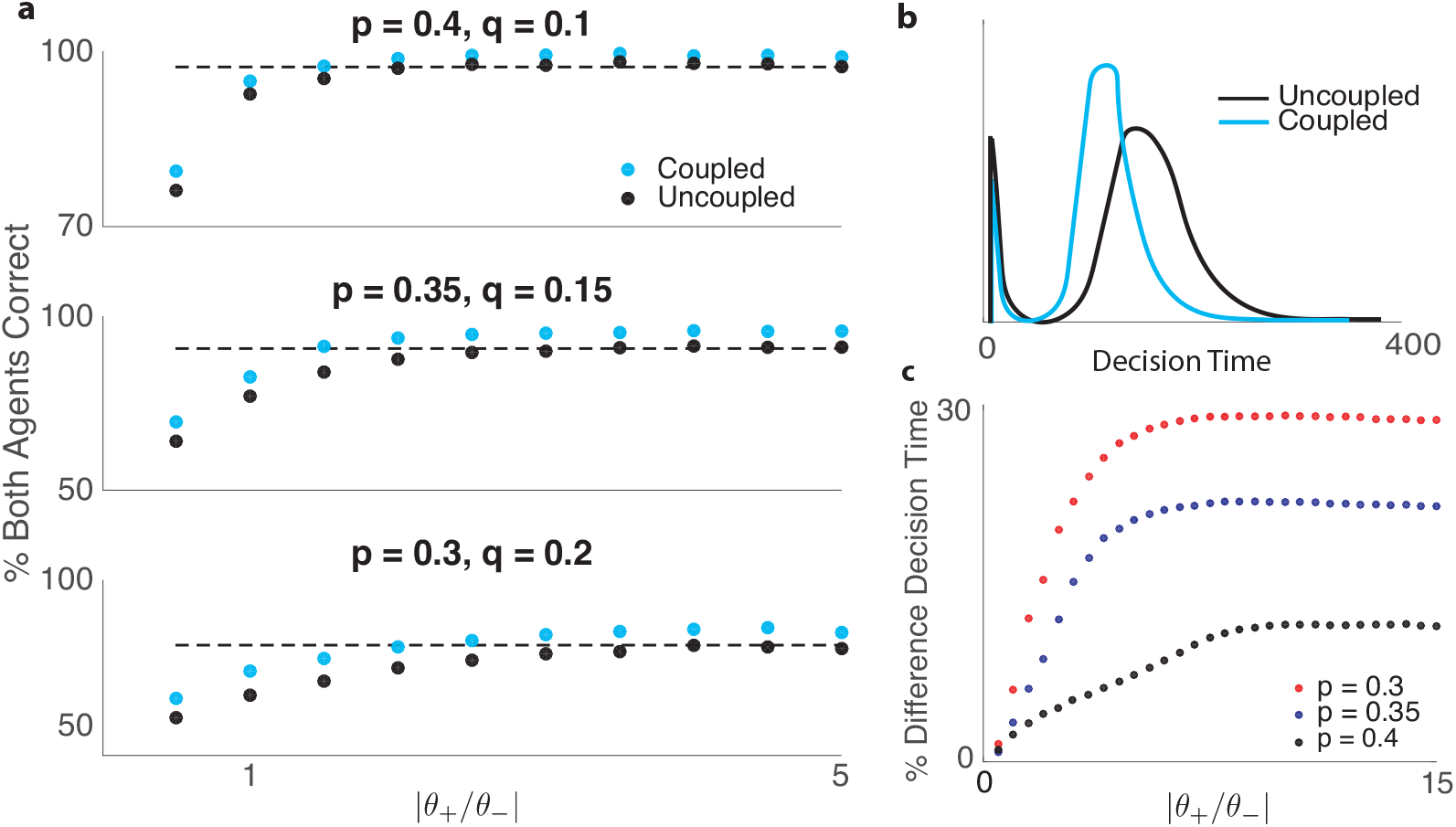
Statistics of decisions in a two-agent network with (coupled), and without (uncoupled) communication from agent 1 to 2. The lower threshold is fixed at θ_−_ = −2 throughout. (a) The fraction of times both agents selected the correct choice as a function of asymmetry in the system as measured by θ_+_/θ_−_. The dashed lines are asymptotic probabilities of the correct choice in the limit θ_+_ → ∞for uncoupled agents. (b) First passage time distributions for the LLR in the case θ_+_ = 40. (c) Relative percent difference in decision times for agent 2 in coupled versus uncoupled trajectories as a function of asymmetry in the system shown for different values of p.

Social information also affects decision times, particularly in the case of strongly asymmetric thresholds (Fig. 4.2b). An early peak in the distributions represents decisions corresponding to the smaller threshold, *θ_−_*, while the latter peak corresponds to the opposite decision when the belief crosses *θ*_+_ ≫ –*θ_−_*. Furthermore, as *p/q* increases, the difference in decision times between the agents decreases (See Fig. 4.2c), as social information speeds up agent 2’s decisions more when private measurements are unreliable. Also, social information plays a more significant role for thresholds *θ_±_* with stronger asymmetry.

**Remark:**

The impact of social information in this example is small, unless the difference in thresholds is very large. However, this impact can be magnified in larger networks: Consider a star network in which an agent observes the decision of *N >* 1 other agents. If these are the only social interactions, the independence of private measurements implies that social information obtained by the central agent is additive. Until a decision is observed, this social information equals 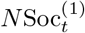, where 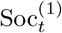 is as defined in Eq. (4.2). Thus the impact of non-decision information can be magnified in larger networks. However, as these cases are computationally more difficult to deal with, we do not discuss them in detail here.

## 5. Two-agent recurrent network

We next assume that the two agents can observe, and react to, each other’s choices. As in the previous section, we assume that at each time the agents synchronously make independent, private observations, 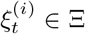, and update their beliefs. The agents then observe each other’s decision state 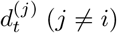, and use this social information to update their belief again. Knowing that a belief has been updated, and observing the resulting decision state can provide new information about an agent’s belief. Hence, social information can continue to be exchanged, even before the next private observation is made (See Fig. 5.1 for illustration).

**Fig. 5.1.**
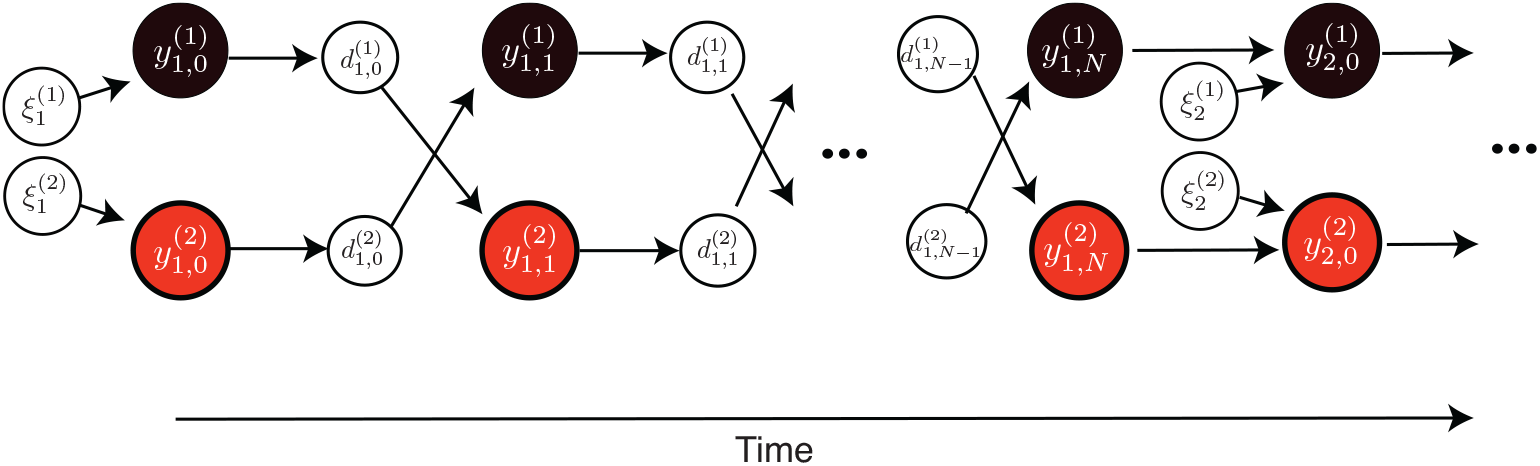
In a recurrent network of two agents the LLRs, 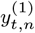 and 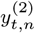, of the two observers are updated recursively. Agents update their belief after private observations, 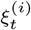, as well as observations of the subsequent series of decision states, 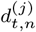, of their neighbor (j ≠ i). After a finite number of steps, Nt, no further information can be obtained by observing each other’s decision, and the two agents make their next private observation, 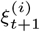, synchronously. The process continues until one of the agents makes a decision.

We describe several properties of this evidence exchange process: As in the case of a unidirectional network, social information is additive. It evolves deterministically up to the time of a decision of the observed agent. Once a decision is made, the social information that is communicated approximately equals the belief threshold (*θ*_+_ or *θ_−_*) crossed by the LLR of the deciding agent. We also show that the exchange of social information after an observation terminates in a finite number of steps either when indecisions provide no further social information, or when one of the agents makes a choice.

Such information exchange has been discussed in the literature on *common knowledge* and rational learning in social networks [33]. This body of work shows that rational agents that repeatedly announce their preferred choice must eventually reach agreement [2, 18, 22, 23, 38]. Typically, it is assumed that information is shared by announcing a preference that can be changed as more information is received. Our assumption that decisions are immutable means that agreement is not guaranteed.

We show how social information is exchanged using induction. We describe the basic case following an observation at *t* = 1 in some detail. Following exchanges are similar, as the belief is updated recursively.

### Social information exchange after the first observation

After the first private observation, 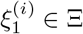, at *t* = 1, the beliefs of the agents are 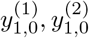, where 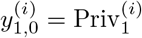. Let

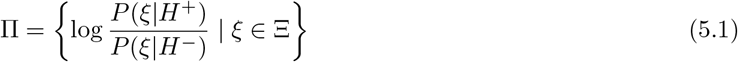

be the set of all possible increments due to a private observation, so that 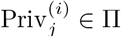, *i* = 1, 2. As the set of possible observations, Ξ, is finite, so is Π. We will describe how the two agents exchange social information with their neighbor until observing a decision, or until no further information can be exchanged. The second subscript, *n* in 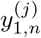, denotes the steps in this subprocess of social information exchange preceding a decision, or next private measurement.

We again associate a belief, 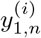, with a corresponding decision state, 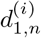, as in Eq. (3.1). If neither of the first two private observations leads to a decision, then 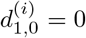, and 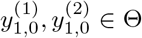, where, Θ *≡* (*θ_−_, θ*_+_) for *i* = 1, 2. Importantly, the fact that agent *i* observed that its counterpart did not decide means that they know 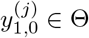 for *i* ≠ *j*

To update their belief, agents compare the probability of all available evidence under the two hypotheses, 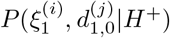 and 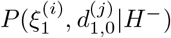. As 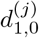 is independent of 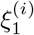 for *i ≠ j*, their updated beliefs are

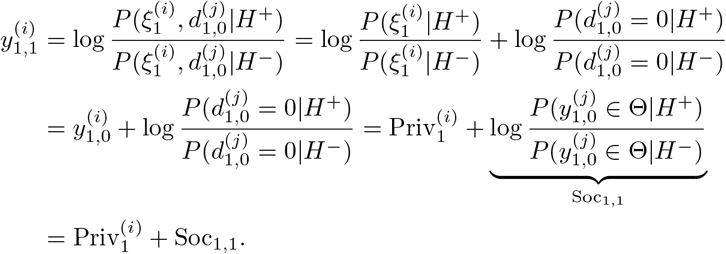

We omit the superscripts on the social information, since 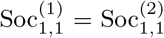. Non-decisions provide the same social evidence to both agents because we assumed the agents are identical. Since the agents know the measurement distributions, *f_±_*(*ξ*), the survival probabilities, 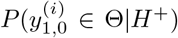, can be computed as in the previous section. Depending on the thresholds and measurement distributions, Soc_1,1_ may be positive, negative, or zero.

If 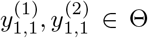, no decision is made after the first exchange of social information, and 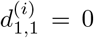 for *i* = 1,2. In this case, agent *i* knows that 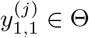 for *j* ≠ *i*, and so

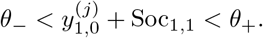

Thus observing 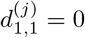 informs agent *i* that 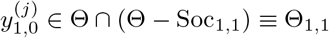 for *i ≠ j*. More precisely,

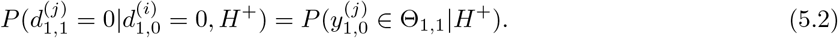

Any initial measurement 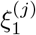 that would lead to a belief 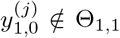 would have lead to a decision at this point. This would end further evidence accumulation. Thus the other agent either observes a decision, or knows that 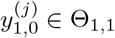.

We will deal with the consequence of observing a decision subsequently, and first consider only observations that do not lead to a decision. For such observations, we can compute the probability of available evidence under either hypothesis by noting that some sequences of decisions have zero probability:

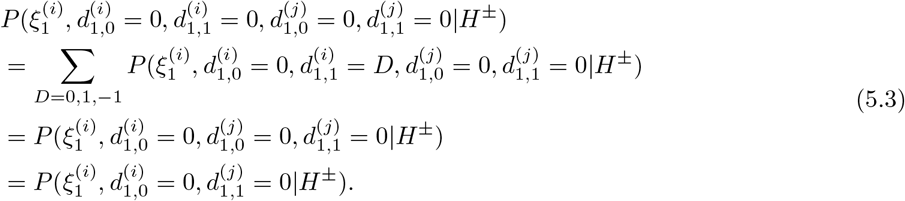

The second equality follows from the fact that an observation 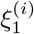 leads to 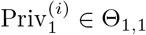, which is inconsistent with 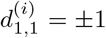. Therefore, 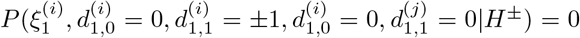. The last equality follows from immutability of decisions.

We also observe that

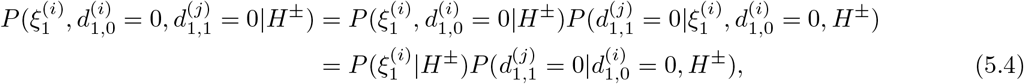

where we used

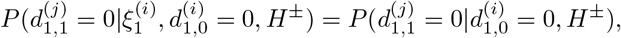

as 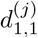 depends only on the observed decision of agent *i*, and 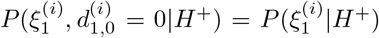, because 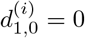 is implied by 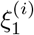 generating 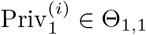.

Combining Eqs. (5.2) and (5.4) gives the updated beliefs,

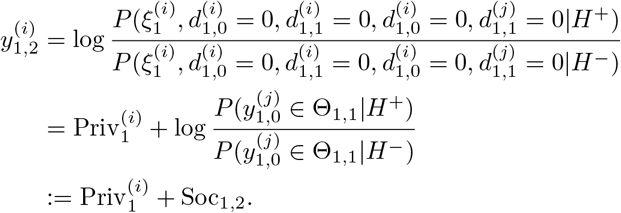

Again, if 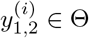, neither agent makes a decision, and both will observe 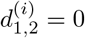.

The same argument used to obtain Eq. (5.3) and Eq. (5.4) shows that for any *l >* 0,

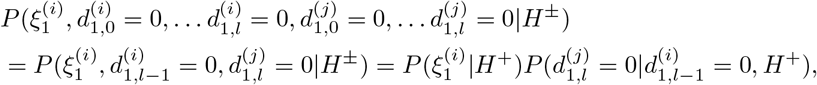

when 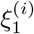 generates 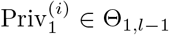, where 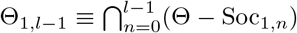. Thus if the *l*-th exchange of social information results in no decision, each agent updates its belief recursively as

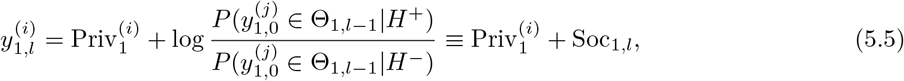

where Soc_1,0_ =0.

This exchange of social information continues until one of the agents makes a choice, or when the absence of a decision does not lead to new social information [23]. The second case occurs at a step, *N*_1_, at which the absence of a decision would not provide any further information about the belief of either agent, that is when

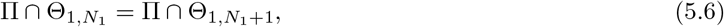

where Π is defined in Eq. (5.1). In this case, 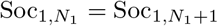, and, if neither agents decide, their beliefs would not change at step *N*_1_ + 1. As both agents know that nothing new is to be learned from observing their neighbor, they then make the next private observation.

We denote the total social information gained after the exchange is complete by Soc_1_, and the belief at the end of this social information exchange by 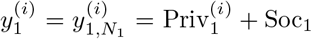. If no decision is made at this point, then agent *j* knows that 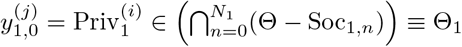

The process can be explained simply: The absence of a decision provides sequentially tighter bounds on the neighbor’s belief. When the agents can conclude that these bounds do not change from one step to the next, the absence of a decision provides no new information, the exchange of social information ends, and both agents make the next private observation.

Importantly, this process is *deterministic*: Until a decision is made, the social information gathered on each step is the same across realizations, *i.e.* independent of the private observations of the agent.

### Social information exchange after an observation at time *t >* 1

The integration of private information from each individual measurement, 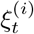, is again followed by an equilibration process. The two agents observe each others’ decision states until nothing further can be learned. To describe this process, we proceed inductively, and extend the definitions introduced in the previous section.

Let 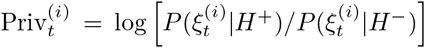 be the private information obtained from an observation at time *t*. For the inductive step assume that each equilibration process after the observation at time *t* ends either in a decision, or allows each agent *i* to conclude that the accumulated private evidence satisfies 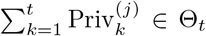 for some set Θ_*t*_ ⊂ Θ. Note that Θ_1_ was defined above, and we will define the other sets in the sequence recursively. We have proved the base step above, since we have shown that following equilibration after observation 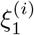, either one of the agents makes a decision, or each agent *i* knows that the private information of its counterpart satisfies 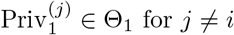 for *j ≠ i*. We then define

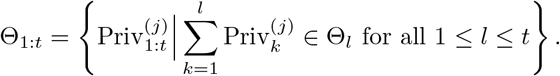

Thus, agent *i ≠ j* knows that any sequence 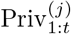 that did not lead to a decision by agent *j* must lie in Θ_1:*t*_, and this is the largest such set. Note that the condition in this definition is given in terms of a sum of LLRs given by private information, so that Θ_1:*t*_is not the same as {Θ_1_*, …* Θ_*t*_}.

To define the social information gathered during equilibration following the observation at time *t*, let Θ_*t,*0_ = Θ_*t−*1_, Soc_*t,*0_ = Soc_*t−*1_, and set

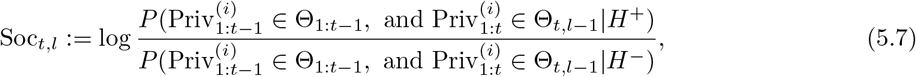

for *l ≥* 1, where 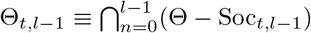.

#### THEOREM 5.1.

*Assume that, in a recurrent network, two agents have made a sequence of private observations, ξ*_1:*t*_, followed by l observations of the subsequent decision states of each other. If neither agent has made a decision, then the belief of each is given by

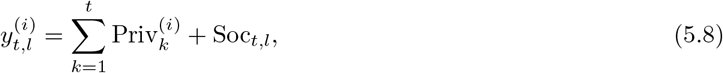

*for* 1 ≤ *t, i* = 1, 2*. The exchange of social information terminates in a finite number of steps after an observation, either when a decision is made, or after no further social information is available at some step l* = *N_t_. The private evidence in* Eq. (5.8) *is a random variable (depends on realization), while the social evidence is independent of realization, and equal for both agents.*

*Proof*. The proof is inductive, with the basic case proved in the previous section. The inductive step follows similarly. By a slight abuse of notation, let 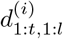 denote the sequence of decision states up to the *l-t* h step in the equilibration process. If no decision has been made at this step, we write 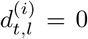. This implies that no previous decision has been made, and we denote this by writing 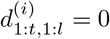.

In the induction step, we assume that equilibration following private observation 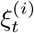 terminates on step *N_t_*. Conditional independence of the observations implies that 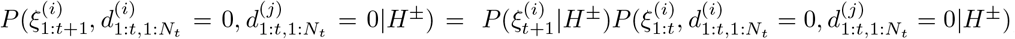, so that

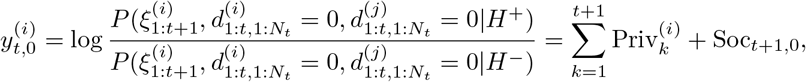

where we used Soc_*t*+1,0_ = Soc_*t*_.

Suppose that no decision has been made in the following *l ≥* 0 equilibration steps, so that 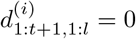 for *i* = 1, 2. For all sequences of measurements 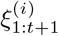 that are consistent with this absence of a decision, we have

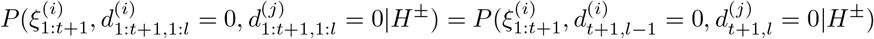

by marginalizing over all intervening decisions, and the final decision of agent *i*, as in Eq. (5.3). As in Eq. (5.4) we have

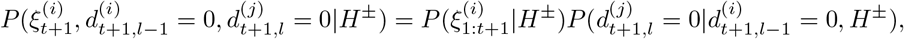

and therefore,

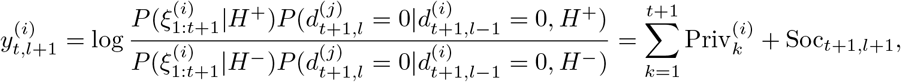

where Soc_*t*+1*,l*+1_ is defined in Eq. (5.7). This exchange of social information stops at step *l* = *N_t_*_+1_ at which point 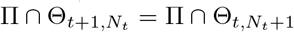, and neither agent learns anything further from the absence of a decision by its counterpart.

### Belief update after a decision

Previously we described social information exchange in the absence of a decision. The following proposition shows what happens when the belief of one of the agents crosses a threshold.

#### PROPOSITION 5.2.

*Suppose that in a recurrent two-agent network, agent i makes a decision after a private observation at time T*^(*i*)^*, during the nth step of the subsequent social information exchange process,* 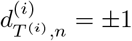*. Then agent j ≠ i, updates its belief as*

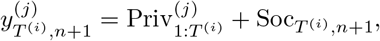

*where*

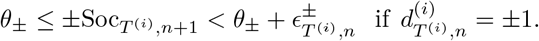

*In the case n* = 0,

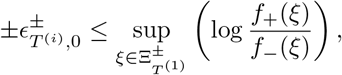

*and* 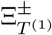 *is the set of all observations leading to a choice of H^±^ at timestep* (*T* ^(*i*)^, 0)*. If n >* 0,

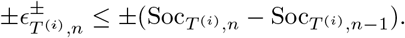

We prove Proposition 5.2 in Appendix D.

As we discussed above, agent *j* may obtain social information before observing the decision of its counterpart. However, this earlier social information is subsumed by the information obtained from observing a decision. If the two agents have different decision thresholds, the expression for the post-decision belief is more involved, but still computable. For simplicity we forgo further discussion of this case.

We next show why this process simplifies when the decision thresholds and evidence distributions are symmetric.

#### PROPOSITION 5.3.

*When the distributions f*_+_ *and f_−_ are symmetric and the agents have the same symmetric thresholds (*±*θ_±_* = *θ), then* Soc_*t*_= 0 *until private evidence leads one of the agents to make a decision. Thus there is no exchange of social information until one of the agents makes a decision.*

*Proof*. The argument is similar to that used to prove Proposition 4.3. We can proceed inductively again: If the two agents have not made a decision after the first observation, by symmetry this does not provide any evidence for either hypothesis *H* = *H^±^*. Observing each other’s decisions after this first observation hence results in no new information,

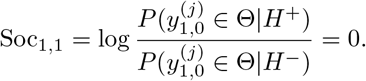

Therefore, the equilibration process terminates immediately, and both agents proceed to the next observation. As shown in Proposition 4.3, further observations provide no new social information, unless a decision is made. The two agents therefore continue to accumulate private information, until one of them makes a decision.

Fig. 5.2 provides examples of evidence accumulation in a two-agent recurrent network. In Fig. 5.2a,b, we illustrate the process with relatively small thresholds and show how the intervals Θ_*n*_shrink at the end of each social evidence exchange (equilibration) following a private observation. Note that the sequence of intervals is the same in both examples because the social information exchange process is deterministic. In this example, equilibration occurs after two steps. In Fig. 5.2c,d, we provide an example with strongly asymmetric decision thresholds. The equilibration of social information between private observations requires several more steps, as shown in the inset of Fig. 5.2c. These examples also illustrate that the beliefs of the two agents do not have to converge, and they do not need to agree on a choice, in contrast to classic studies of *common knowledge* [2, 22, 33].

**Fig. 5.2.**
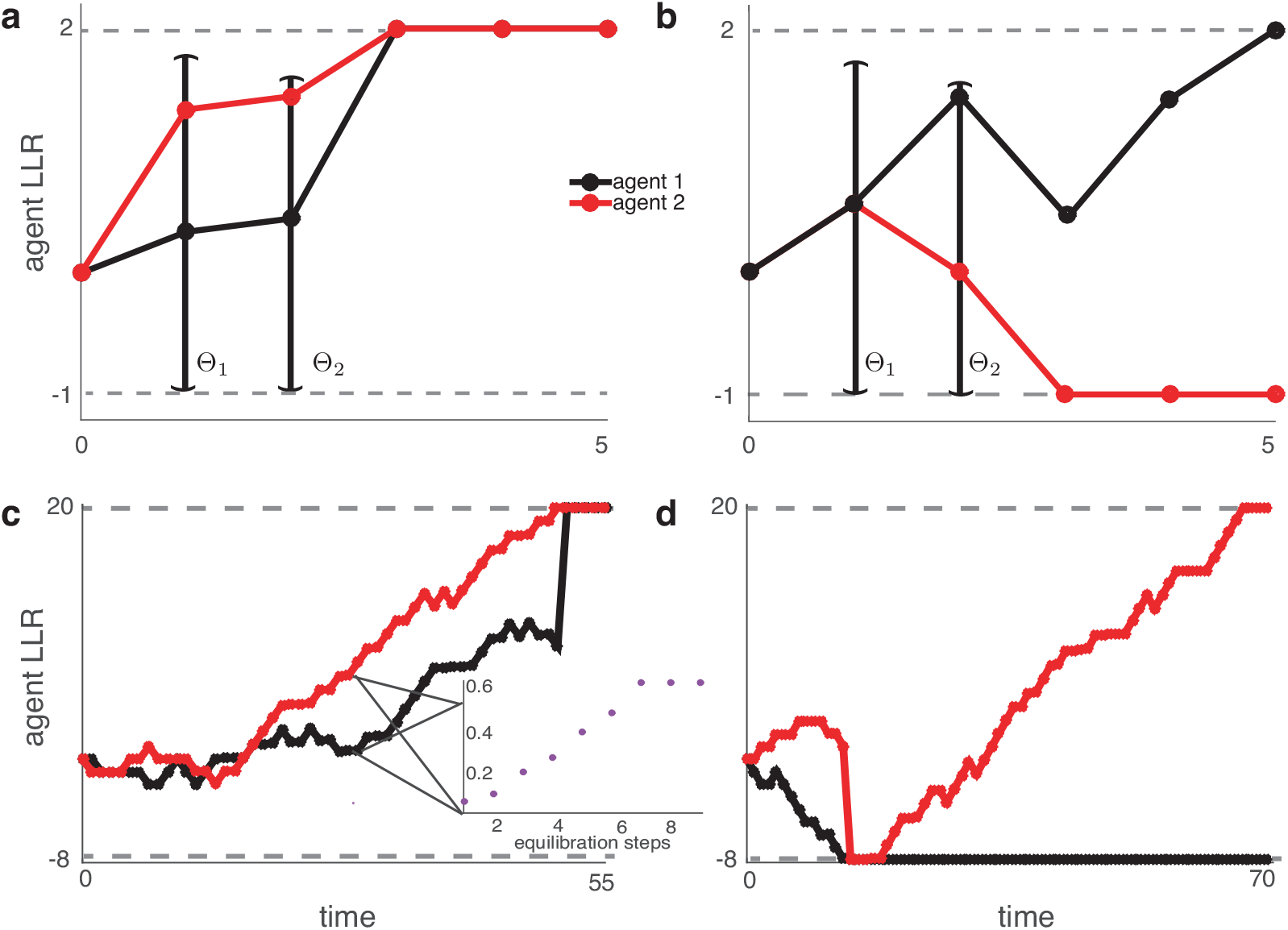
The belief (LLR) of agents 1 and 2 in a recurrent network with asymmetric thresholds when (a) the agents agree, and (b) agents disagree on a final choice. Also shown are the intervals Θ_1_, Θ_2_, resulting from the equilibration following the first two observations. Each agent knows that the LLR of its counterpart lies in this interval after equilibration ends. Although the beliefs evolve stochastically, the sequence Θ_1_, Θ_2_ is fixed across realizations. Here, p = e/7, q = 1/7, and s is determined from s = 1 p q. Also, θ_+_ = 2 for (a)-(b) and θ_+_ = 20 for (c) - (d). θ_−_ = 1 for (a)-(b) and θ_−_ = 8 for (c)-(d). Inset: Social information obtained from equilibration converges in seven steps at the indicated time.

## 6. Accumulation of Evidence on General Networks

In networks that are not fully connected, rational agents need to take into account the impact of the decisions of agents that they do not observe directly. To do so they marginalize over all unobserved decision states. This computation can be complex, even when thresholds and evidence distributions are symmetric.

To illustrate we begin by describing the example of agents with symmetric decision thresholds and measurement distributions on a directed chain. Symmetry makes the computations more transparent, as the absence of a decision is not informative about the belief of any agent, hidden or visible. Social information is therefore only communicated when an agent directly observes a choice. Such an observation leads to a jump in the agent’s belief, and can initiate a cascade of decisions down the chain [10].

We note that once an agent in the network makes a decision, symmetry can be broken: Agents know that all others who have observed the decision have evidence favoring the observed choice. As we have seen, once symmetry is broken even the absence of a choice can provide social information. In this case rational agents must equilibriate their beliefs using known survival probabilities, as discussed in the previous section. We do not analyze this case.

### 6.1. Terminology and Notation

In a social network of *N* agents, we again assume that each agent makes a private observation at every time step. After incorporating the evidence from this observation the agent then updates its decision state and shares it with its neighbors. A directed edge from agent *j* to *i*, denoted by *j* → *i*, means that agent *j* communicates its decision state to agent *i*. The set of neighbors that agent *i* observes is denoted by *N* ^(*i*)^ = {*j* : *j* → *i*}.

Agent *i* thus receives social information from all agents in *N*^(*i*)^. However, agent *i* also needs to take into account decisions of *unobserved* agents. We define the set of all agents not visible to agent *i* as

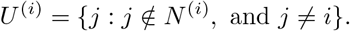

Thus all agents in the network, besides agent *i*, belong to *N*^(*i*)^ or *U*^(*i*)^. Let *W* = {1, 2, 3*, …, N*}. Then *W* = *N*^(*i*)^ ∪ *U*^(*i*)^ ∪ {*i*} and *N*^(*i*)^ *∩ U*^(*i*)^*, N*^(*i*)^ *∩* {*i*}*, U*^(*i*)^ *∩* {*i*} all equal ∅.

We denote the set of decisions by the neighbors of agent *i* following the observation at time *t* by 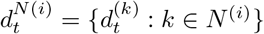. Similarly, the set of the decisions by unobserved agents is denoted by 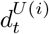. More generally, 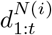 denotes the sequence of decision states of the neighbors of agent *i* up to and including the decision following the observation at time 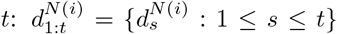. We will see that in the case of symmetric thresholds and observations no equilibration occurs, so these decision states describe information available to each agent in the network completely until one of the agents makes a choice. At time *t*, the private and social observations obtained by agent *i* are therefore 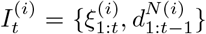. As before, we denote the private information by 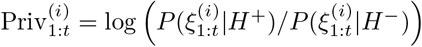.

### 6.2. Non-decisions

We next show that in social networks of agents with symmetric thresholds and measurement distributions, the agent’s beliefs have two properties that simplify computations: An absence of a decision is uninformative, and once a decision is observed, the resulting social information is additive.

#### PROPOSITION 6.1.

*Assume all agents have symmetric thresholds and evidence distributions. If neither agent i, nor any of its neighbors in N*^(*i*)^ *have made a decision by time t, then the belief of agent i is*

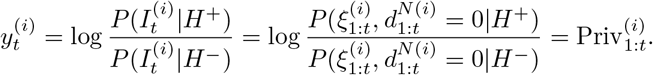

*Thus social information is only obtained from observing a choice.*

*Proof*. The claim follows from an argument similar to that given in Sections 4 and 5. The main difference is that agent *i* here marginalizes over unobserved decision states.

By definition, the only direct social information agent *i* has is the set of decision states of its neighbors 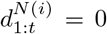 to time *t*. First we can split the private evidence from the social evidence conditioned on the agent’s observations,

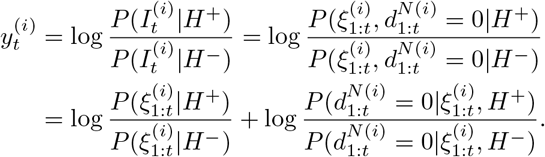

We therefore need to show that

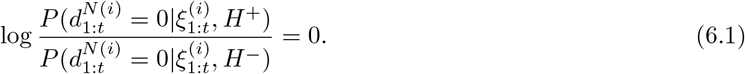

Marginalizing over all possible combinations of decision states for unseen neighbors, 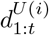, and observations of neighbors, 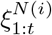, yields

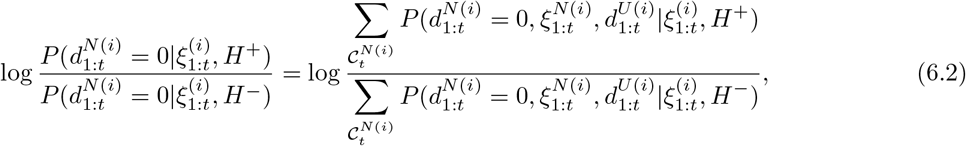

where 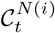 denotes all possible chains 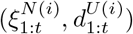 of social and private observations that neighbors of agent *i* could have made without reaching a decision by time *t*. For a decision history of the unobserved agents 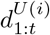 let 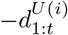, be the opposite decision history, flipping the sign of each decision in the vector 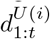, and leaving non-decisions unaffected. The vector of zeroes equals its negation. Likewise, for a private observation history, 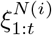, of the neighbors, let 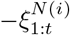 be the opposite sequence of observations. Symmetry in thresholds and measurement distributions implies 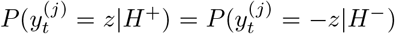 for all *z* ∈ Ξ. By symmetry,

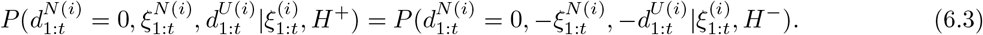

Therefore in Eq. (6.2), for every chain 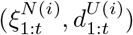 appearing in the numerator, its negation 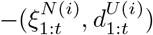 also appears in the denominator. The sums in the numerator and denominator on the right hand side of Eq. (6.2) are equal, and hence Eq. (6.1) holds.

It remains to show that none of the agents engage in equilibration. Note that after its first private measurement, no agent *i* obtains further information from a non-decision of its neighbors. The equilibration process therefore does not start. The same holds after further private measurements by induction.

Thus, before decisions are made, agents independently accumulate evidence. Decision states are only informative when an agent makes a choice. As in the case of two agents, a choice by any agent will lead to a jump in the belief of all observing neighbors.

**Remark:** Suppose that two neighboring agents, *j* and *k*, both observe an agent making a choice. Agent *k* now knows that the belief of agent *j* must have increased by a known amount due to the mutually observed decision. Agent *k* thus gains knowledge about the belief of agent *j*, even if *j* does not immediately decide. The symmetry is therefore broken, and henceforth the situation is similar to the asymmetric case discussed in Section 5. This introduces additional drift, and potentially the need for equilibration, until agent *k* or *j* make decisions. We thus concentrate on the dynamics up to the first decision, but discuss the general case in more detail in some of our examples.

### 6.3. Example of Marginalization: 3-Agents on a Line

We demonstrate the computations needed to account for unobserved agents using the example shown in Fig. 6.1. In this case a decision by agent 3 is not observed by any other agent. If agent 1 makes a decision first, then agent 2 continues to accumulate information as in the case of two unidirectionally coupled agents discussed in Section 3. It is therefore sufficient to only consider the case when agent 2 makes a decision first at time *T*^(2)^.

**Fig. 6.1.**
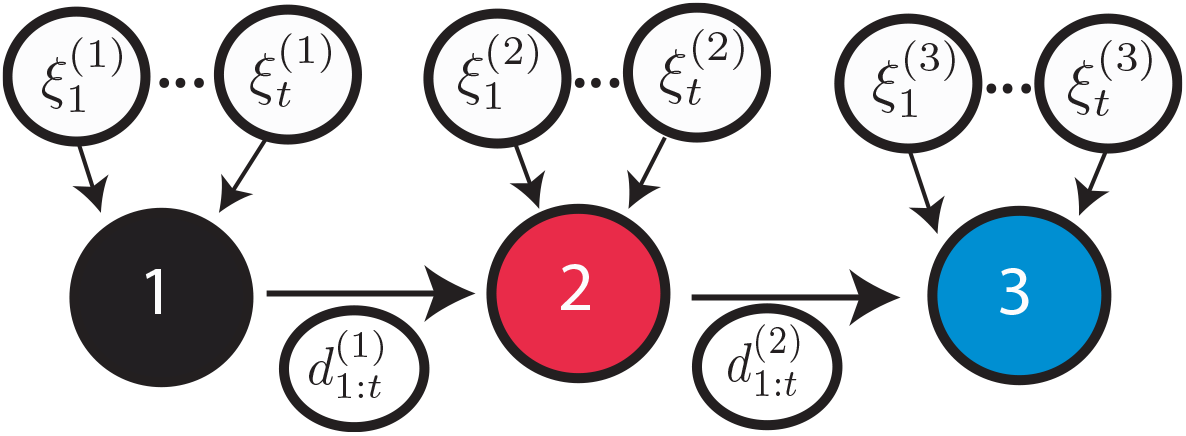
Agents on a line. Agent 3 observes the decisions of agent 2, but not of agent 1. However, agent 3 still needs to take into account the possibility that a choice by agent 1 has caused a decision by agent 2.

As shown in Proposition 6.1, agent 3 has no social information before observing a decision by agent 2. After observing a decision by agent 2, agent 3 updates its belief by marginalizing over possible decision histories of agent 1:

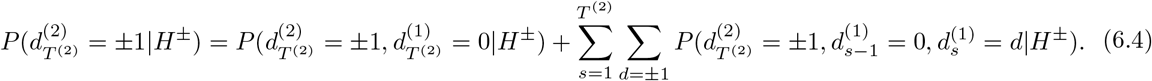

Intuitively, a choice of agent 2 can be triggered by either: (a) A private observation leading to the belief 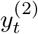 reaching one of the thresholds, *θ_±_*. This possibility is captured by all terms in Eq. (6.4) except

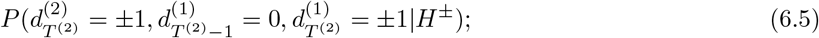

or (b) The decision of agent 1 causing a jump in the belief 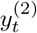 above threshold *θ*_+_ or below threshold *θ_−_*. This second possibility is captured by the term in Eq. (6.5). An argument equivalent to that in Proposition 4.1 shows that the social information communicated in the first case is close to *±θ* for 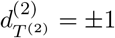. However, in the second case the belief of agent 2 at the time of decision is in the range 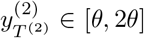 for 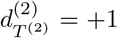 or 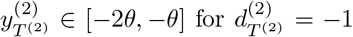, modulo a small correction. Agent 3 updates its belief by weighting both possibilities, and hence increases its belief by an amount greater than *θ*. We provide details of this argument in Appendix E, and show that the excess increase in belief of agent 2 can be large, even when the increments in private information, 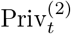, are small.

In Fig. 6.2, we illustrate how including social information affects key metrics in decision-making: decision-time and accuracy. When *p/q* is not too large, and hence the signal-to-noise ratio (SNR) of observations is low, the exchange of social information impacts both the accuracy and response time of agents significantly. Agents in a social network choose the correct decision more often compared to isolated agents, as the impact of social information is significant. When *p/q*, and hence SNR, is large, including social information does not benefit the agents strongly: accuracy and decision time of agents in a network are similar to isolated agents. Private information dominates the decision making process in these cases.

**Fig. 6.2.**
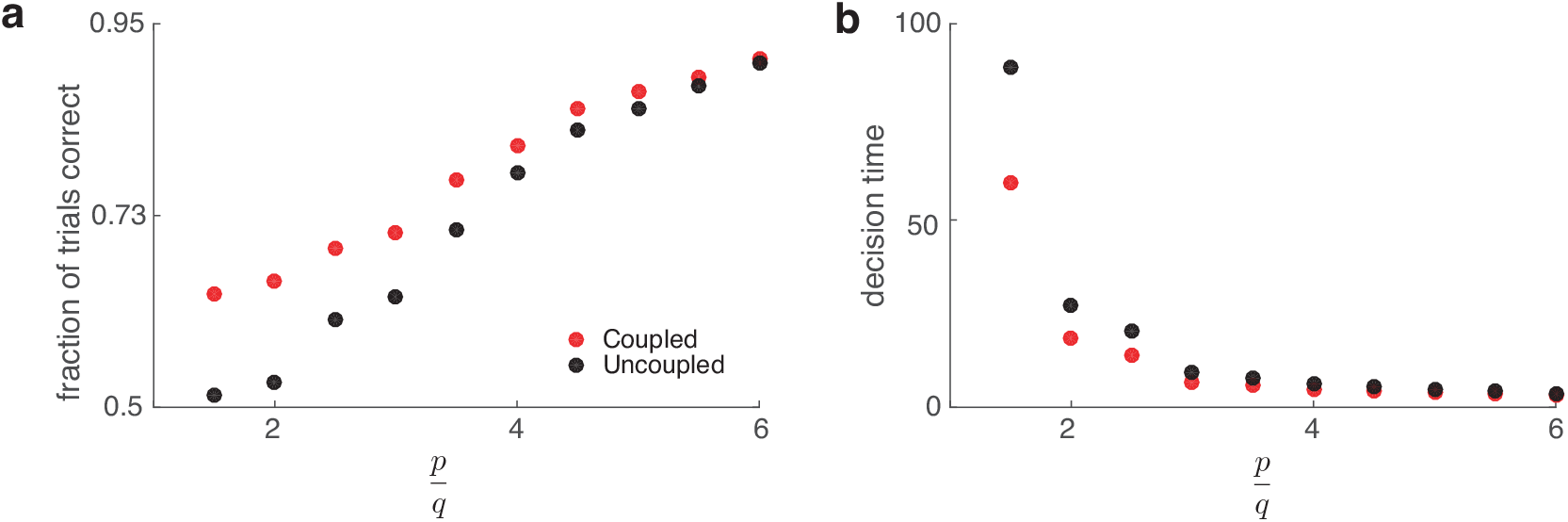
The performance of the three agents on a directed line is better than that of independent agents. (a) The fraction of trials for which all three agents make the correct choice. This quantity is larger when agents are allowed to exchange information (coupled), than when agents make decisions independently (uncoupled). (b) Time required for all three agents to make a decision. Average decision times are also smaller when agents exchange information. Here, |θ_±_|= 30, and ratio p/q determines the noisiness of the measurements, as described in Appendix C.

Marginalization becomes more complex when the unseen component, *U* ^(*i*)^, of the network is larger. For instance, if we consider a long, directed line of *n* agents, when the *k*-th agent makes a decision, the *k* + 1-th agent must marginalize over the preceding *k –* 1 agents. If the resulting jump in the belief, *y*_t_^(*k*+1)^, exceeds 2*θ*, this triggers an immediate decision in agent *k* + 1, and all successive agents. This is equivalent to herding behavior described in the economics literature [1, 5].

## 7. Three-Agent Cliques

In cliques, or all-to-all coupled networks, all agents can observe each others’ decisions, and *U* (*i*) = for all *i*. This simplifies the analysis, as no agent needs to marginalize over the decision states of unobserved agents. We start by discussing the case of three-agent cliques in some detail, and proceed to cliques of arbitrary size in the next section.

As we are assuming symmetry, social evidence is shared only after the first agent makes a choice. Without loss of generality, we assume that this was agent 3, and that 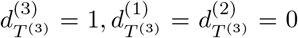. There is a small probability that private information leads to a concurrent decision by multiple agents, and we describe this case at the end of the section. The social information obtained from a decision by agent 3 may or may not drive agent 1 or 2 to a decision. Both the presence and absence of a decision by either agent reveals further social information to its undecided counterpart, and we assume that all undecided agents wait until no further social information can be obtained before gathering further private information. We will see that, akin to the equilibration process, there can be a number of decision-making rounds before evidence accumulation continues.

There are three possible outcomes following the decision of agent 3 (See Fig. 7.1): If 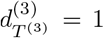 and 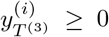, for *i* = 1, 2, then both remaining agents decide immediately, 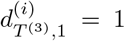, and the evidence accumulation process stops. We therefore only examine cases where (i) 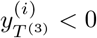 for *i* = 1 or *i* = 2 (but not both), and (ii) 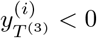 for both *i* = 1, 2.

**Fig. 7.1.**
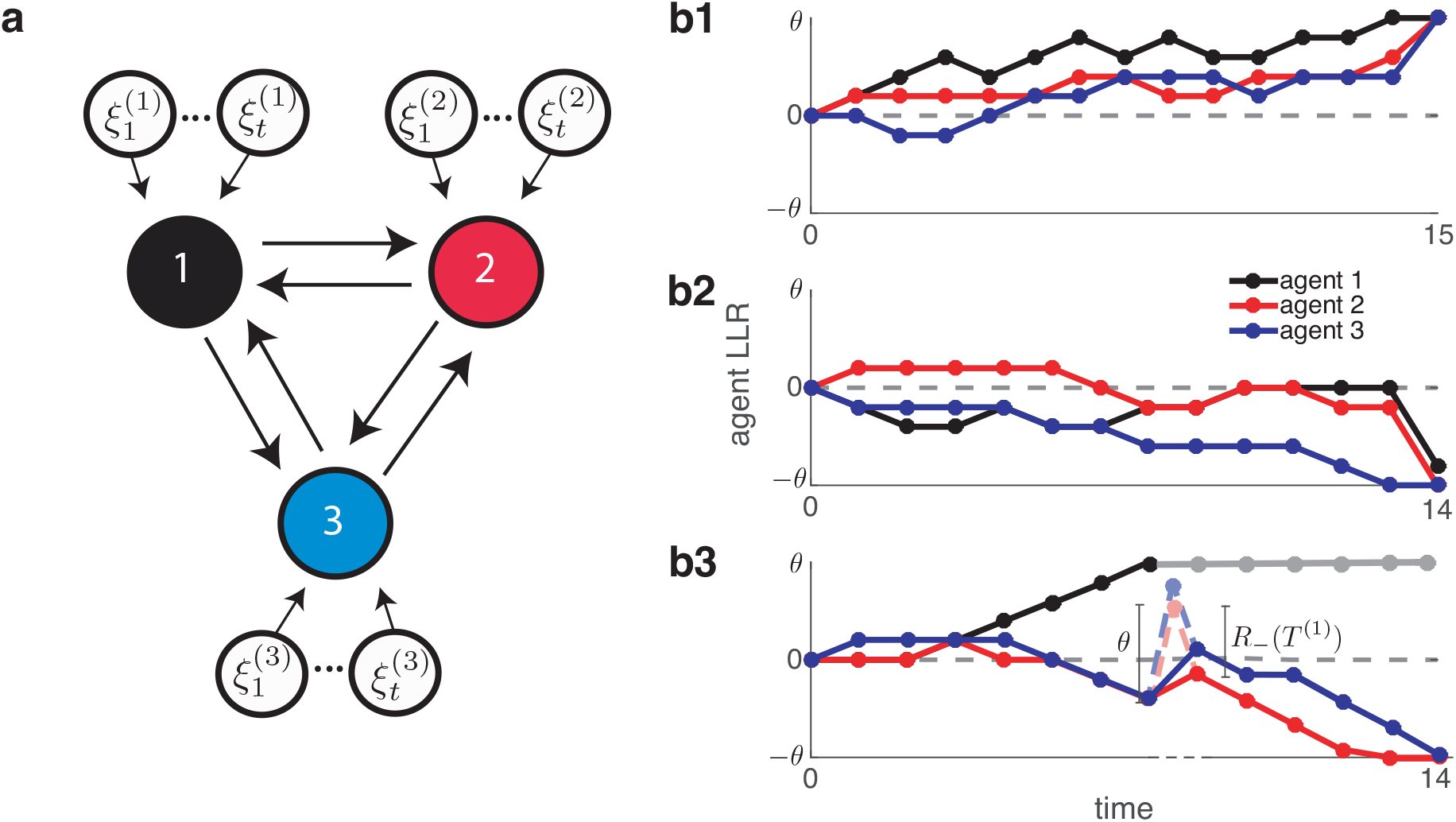
(a) In a three-agent clique all three agents make independent observations, and observe each other’s decisions. (b) Three main possibilities follow the decision of agent 3: (b1) If the beliefs of the undecided agents are both positive, both follow the decision of agent 3; (b2) If the decision causes only one of the remaining agents to decide (agent 2 here). This secondary decision leads to a further update in the belief of the remaining agent equal to R_−_(T ^(3)^) (agent 1 here); (b3) If neither of the remaining agents decides, they observe each other’s indecision, and update their belief by R_−_(T^(1)^) < 0. This update cannot lead to a decision, and both agents continue to accumulate private evidence. The dashed portion of the belief trajectory shows the intermediate steps in social information exchange: Each agent’s belief consists of information due to its own private observation, a jump θ > 0, followed by a jump R_−_(T^(1)^) < 0. No decision can be reached, and the two agents continue to accumulate information, but now have further knowledge about each other’s belief. Cases in which the private evidence leads to a simultaneous decision by two or three agents are not shown.

As in the equilibration process described previously, agents update their beliefs based on the observed decision state of their counterpart until no further information can be gained from the process. Before observing the decision of agent 3, and after the private observation at time *T*^(3)^, the beliefs of agents *i* = 1, 2 are 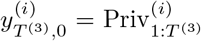 by Proposition 6.1. They next account for the decision 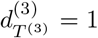 that reaches both simultaneously:

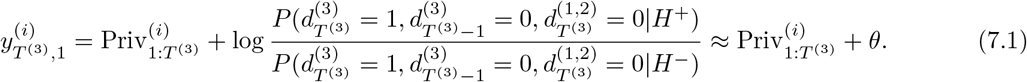

The social information obtained from agent 3’s decision should incorporate the increment 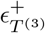 discussed in

### Proposition 4.1.

For a clearer exposition, we assume private evidence is weak, and 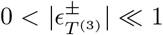, and we thus use the approximation 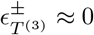 from here on.

Any agent who remains undecided after the update given by Eq. (7.1) will update their belief iteratively in response to the decision information of their neighbor. We describe this process in the cases in which at least one agent remains undecided after observing the decision of agent 3.

*Case (i) - One Agent Undecided.* Without loss of generality, we assume 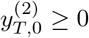 so that 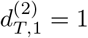. After observing the decision of agent 3, the belief of agent 1 is 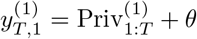. After observing the decision of agent 2, a straightforward computation gives:

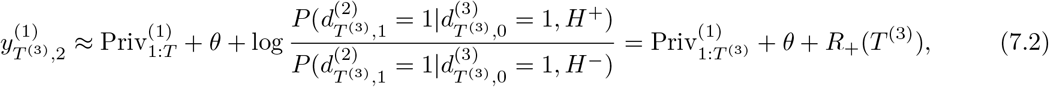

Where

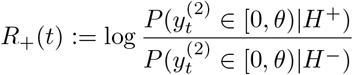

since agent 2’s belief must have been non-negative before observing the decision of agent 3. We prove in Proposition F.1 that *R*_+_(*t*) *< θ*. However, this increment in belief may be sufficient to lead to a decision by agent 1. If not, agent 1 continues to accumulate private evidence on the next time step.

We will estimate *R*_+_(*t*) in arbitrarily large cliques in section 8. In particular, *R*_+_(*t*) may be computed explicitly in the same way as *S_±_*(*t*) (See Appendix C).

*Case (ii): Two Agents Undecided.* If 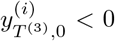 for *i* = 1, 2, then both agents remain undecided upon observing a decision by agent 3. After each observes this absence of a decision in its counterpart, it follows from the computations that lead to Eq. (7.2), that each updates its belief as

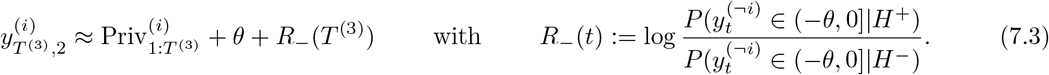

Due to symmetry, the new social information is equal for both agents.

Note that *R_−_*(*t*) *≤* 0 and also *|R_−_|*(*t*) *< θ*, as shown in Proposition F.1. Therefore Eq. (7.3) shows that this second belief update cannot lead to a decision, and 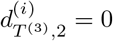 for *i* = 1, 2. After this second step, both agents, *i* = 1, 2, know that

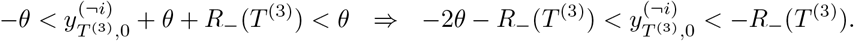

Since *R_−_*(*T* ^(3)^) ∈ (–*θ,* 0], this does not reveal any additional information, and the exchange of social information stops.

At the end of this exchange, both remaining observers know that the belief of the other is in the non-symmetric interval, 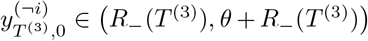. Therefore, future non-decisions become informative, and equilibration follows each private observation as in the case of asymmetric decision thresholds discussed in Section 5.

### Concurrent Decisions

If the first decision is made simultaneously by two agents, the remaining agent receives independent social information from both. When the two deciding agents disagree, the social information they provide cancels. If the two agents agree on *H^±^*, the undecided agent increases its belief by ±2*θ* and follows the decision of the other two.

The exchange of social information increases the probability that all three agents reach a correct decision (Fig. 7.2a), and decreases the time to a decision, both of a single agent, and all agents in the clique (See Fig. 7.2b). The addition of strong pulsatile increments of social information leads to many trials in which agents’ beliefs attain values well beyond the decision thresholds ± *θ*, mapping to accuracy that is higher than would be obtained by beliefs at or slightly beyond threshold. This is particularly pronounced when the SNR of private measurements is low, and uncoupled agents’ final beliefs are not pushed much beyond their decision thresholds. When private observations are more reliable, social information becomes less important.

**Fig. 7.2.**
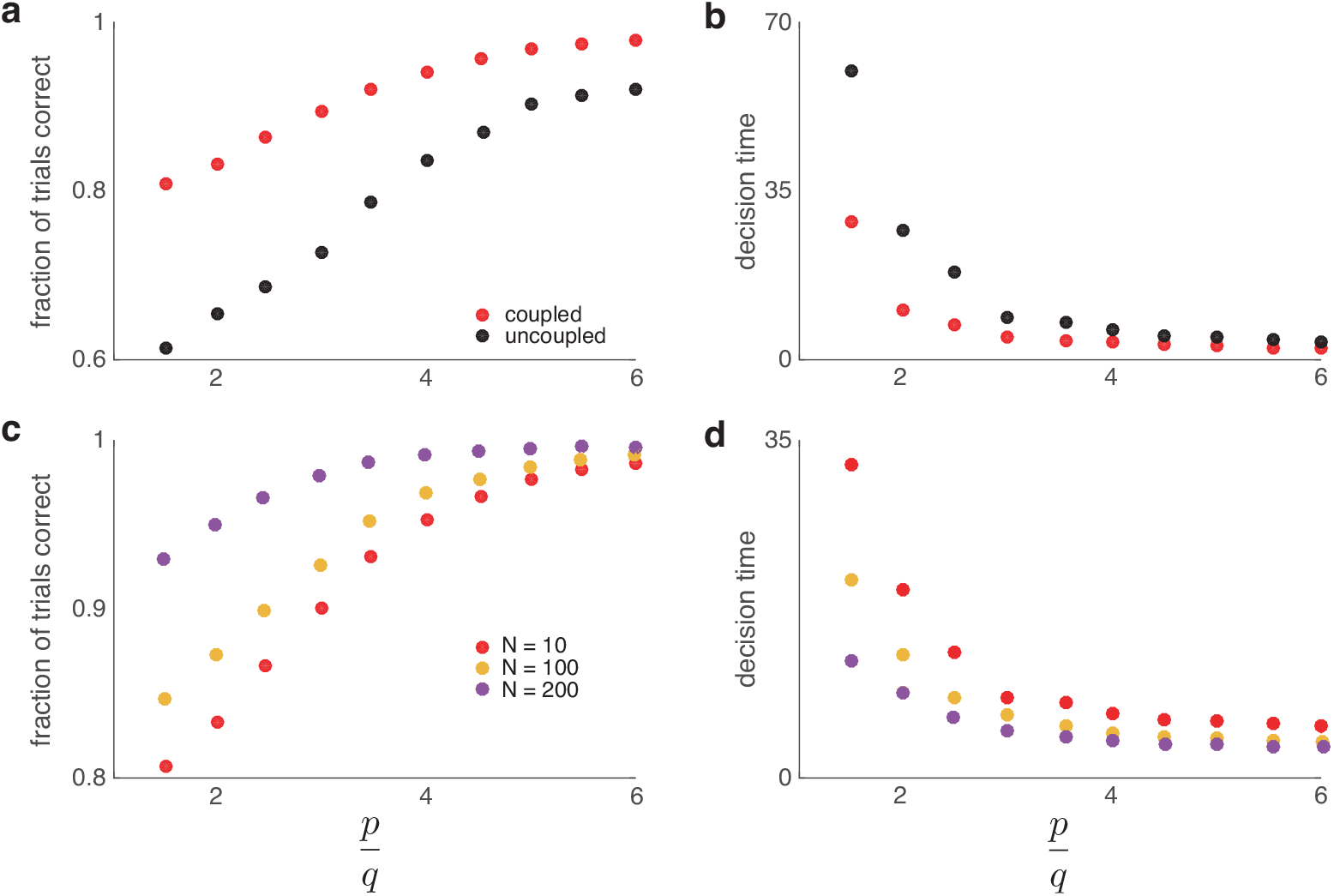
(a) In a three-agent clique the probability that all three agents make the correct choice is larger when social information is exchanged. (b) The average time it takes all three agents decide is smaller when the agents communicate, since social information pushes agents closer to threshold. Both effects are more pronounced when the SNR of private evidence is low, i.e p/q is small. (c) As the number of agents in a clique, N, increases, the probability that all agents in the clique make the correct choice grows. The difference is more pronounced when SNR is low. (d) Larger cliques provide more social information reducing the average decision times. Here, |θ_±_ |= 30, and p, q are defined as in Section 4 and Appendix C. These simulations do not account for the situations where equilibration after the first decision is necessary.

In the next section, we will describe similar cascades of decisions in larger cliques. Note that as we increase the size of the network, the increase in accuracy and decrease in decision time due to the inclusion of social information both grow (See Fig. 7.2c,d). We next discuss the computations following a choice in such larger cliques.

## 8. Larger Cliques

We consider a clique of *N* agents who all have identical symmetric decision thresholds, and measurement distributions. No social information is exchanged until one or more agents makes a choice at time *T*. In what follows, we focus on the case of a single agent making the first decision. The case in which two or more agents simultaneously make the first decision can be easily analyzed as in the three agent clique, and it often leads to all agents deciding subsequently, so we do not discuss it further.

Without loss of generality, we assume agent 1 makes the first decision, and chooses *H*^+^. This means 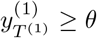, and thus every other agent, *i*, updates their belief to

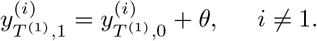

As earlier, we assume that the excess information, 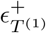 (See Proposition 4.1 for definition), provided by a decision is negligible.

Upon observing a decision, the remaining agents stop making private observations and exchange social information until no further social information is available. Observing 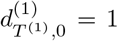 leads any agent *i* with belief 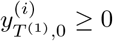 to the same decision. We denote the set of these agents by *A*_1_, and call these the *agreeing agents*. We will see that there can be multiple waves of agreeing agents. The agents whose beliefs satisfy 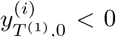 update their belief to 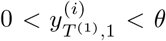, but do not make a decision. We denote the set of these agents by *U*_1_, and call these *undecided agents*.

Once agent 1 chooses *H*^+^ at time *T* ^(1)^, the set of decisions from the first wave of agreeing agents follows at the social information exchange step, (*T*^(1)^, 1). These decisions are conditionally independent, given the observed decision of agent 1. Thus, each agreeing agent independently provides additional evidence for *H*^+^, while each undecided agent provides evidence for *H^−^*. As in the case of three agents, the social information provided by an agreeing agent is *R*_+_(*T* ^(1)^), and for a disagreeing agent is *R_−_*(*T* ^(1)^) = − *R*_+_(*T* ^(1)^), where *R*_+_(*T* ^(1)^) is given in Eq. (E.3). The equality follows from our assumption of symmetry: See Appendix G for a proof. Note that *N* = *a*_1_ + *u*_1_ + 1, where *a*_1_ and *u*_1_ are the number of agreeing and undecided agents in the first wave following the decision of agent 1 at time *T* ^(1)^. All undecided agents thus update their belief as

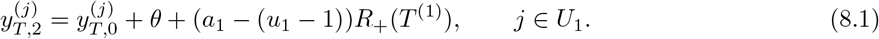

Note, each undecided agent observes all agreeing agents, *a*_1_, and all undecided agents but itself.

We will see that rounds of social evidence exchange can follow. Let

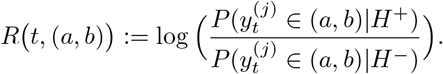

Note that *|R t,* (*a, b*) *| ≤ θ* by Proposition F.1, and that *R t,* [0*, θ*) = *R*_+_(*t*), *R t,* (*−θ,* 0] = *R_−_*(*t*).

Each remaining undecided agent has now observed the decision of the *a*_1_ agreeing agents, and the indecision of the other *u*_1_ –1 undecided agents besides itself. Eq. (8.1) implies several possibilities for what these agents do next:

1. If the number of agreeing agents, *a*_1_, exceeds the number of undecided agents, *u*_1_, by a sufficient amount, (*a*_1_–(*u*_1–_ 1))*R*_+_(*T* ^(1)^) ≥2*θ*, then all the remaining undecided agents, *j∈U*_1_, go along with the decision of the first agent, and choose *H*^+^. Thus the second wave of agreeing agents encompasses the remainder of the network.
2. If the number of undecided agents, *u*_1_, exceeds the number of agreeing agents, *a*_1_, by a sufficient amount, (*a*_1_ *−*(*u*_1_ *−*1))*R*_+_(*T* ^(1)^) *≤ −*2*θ*, then all undecided agents update their belief to 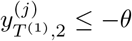*, j*∈ *U*_1_. This is a somewhat counterintuitive situation: If sufficiently many agents remain undecided after observing the first agents’ choice, then, after observing each other’s absence of a decision, they all agree on the opposite. Thus a wave of agreement is followed by a larger wave of disagreement.
3. If –2*θ <* (*a*_1_–(*u*_1_–1))*R*_+_(*T* ^(1)^) *< – θ*, then some of the remaining agents may disagree with the original decision and choose *H^−^*, while others may remain undecided. We call the set of disagreeing (contrary) agents *C*_2_, and the set of still undecided agents *U*_2_, and denote the sizes of these sets by *c*_2_, and *u*_2_ respectively. Note that *U*_1_ = *C*_2_ ∪ *U*_2_. The agents in *U*_2_ know that 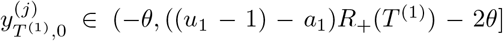 for all *j* ∈ *C*_2_, and 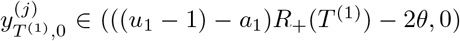 for all *j* ∈ *U*_2_. All agents in *U*_2_ thus update their belief again to

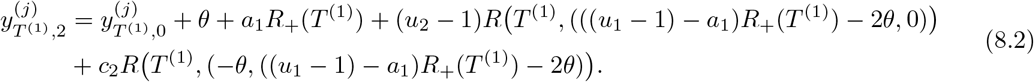 This update includes the social information obtained from the initial observation, *θ*, and from the agreeing agents in the first round, *a*_1_*R*_+_(*T* ^(1)^). The last two summands in Eq. (8.2) give a refinement of the social information from the originally undecided agents in *U*_1_. As a result of this second update some undecided agents in *U*_2_ can make a choice. If so, the process repeats, until no undecided agents are left, or no decisions occur after an update. This process thus must terminate after a finite number of steps. This is akin to the equilibration of social information described earlier, but involves observed choices and occurs across the network. Importantly, the process is realization dependent, as it depends on the private evidence accumulated by the undecided agents. After the process is complete, symmetry is broken, and social information equilibration occurs after each further private measurement.
4. If –*θ* ≤ (*a*_1_ –(*u*_1_–1))*R*_+_(*T* ^(1)^) ≤0, then no agents in *U*_1_ make a decision, and no undecided agent obtains any further social information. They thus continue to gather private information. Symmetry is broken as they know that 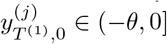 ∈ (–*θ*,0]for all remaining agents, *j* ∈ *U*_1_.
5. If 0 *<* (*a*_1_ *−* (*u*_1_ *−* 1))*R*_+_(*T* ^(1)^) *< θ*, some undecided agents may choose *H*^+^, and some may remain undecided. We call the first set *A*_2_ and the second *U*_2_, and denote the sets’ cardinality by *a*_2_ and *u*_2_, respectively. In this case, *U*_1_ = *A*_2_ ∪ *U*_2_. All undecided agents *j* ∈ *U*_2_ know that 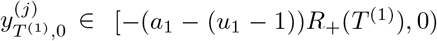 for all *j* ∈ *A*_2_, and 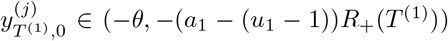for all *j* ∈ *U*_2_. They thus update their belief to

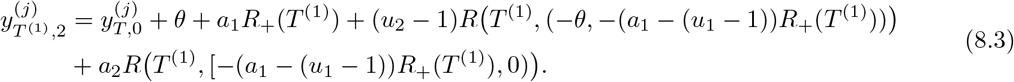 As a result of this update, some agents may make a choice. If so, the process continues as discussed in 3. To summarize, before any agent in a clique makes a decision, each agent accumulates private evidence independently. A given agent will make the first decision, say *H*^+^. All other agents with positive evidence will follow this choice in a first wave of agreement. Undecided agents modify their evidence according to Eq. (8.1). How many agents agree *a_n_*, disagree *c_n_*, or remain undecided *u_n_* in exchange *n* depends on the numbers of agreeing, disagreeing, and undecided agents from previous steps: *a*_1:*n−*1_, *c*_1:*n−*1_, and *u*_1:*n−*1_.

### Large System Size Limit

Let *α_N_* be the probability that the first agent to decide in an *N*-agent clique makes the correct decision. We conjecture that as the clique size *N*→∞, the probability that the majority of the agents make the correct decision goes to 1; furthermore, all agents will make the correct decision with probability *α_N_* . An asymptotic argument for this result in the special case of a biased random walk on an integer lattice is straightforward. However, a detailed proof of the general claim would be lengthy, so we save a thorough treatment of this problem for future work.

In Fig. 8.1 we plot information about the behavior of cliques of various sizes, hinting at trends that emerge as *N* →∞ . We assume the correct state is *H*^+^. As the size of the clique grows, so does the amount of information available to undecided agents after the first wave of decisions, *i.e.* the members of the set *U*_1_ (See Fig. 8.1b). As clique size grows, the first decision occurs earlier (Figs. 8.1c-d). In particular, as *N* grows, the first decision time approaches *θ/* log(*p/q*)–the minimum number of steps required to make a decision. Moreover, most agents accrue evidence in favor of the correct decision at an earlier time. By the time the first decision occurs, the majority of the clique will be inclined toward the correct decision. The ensuing first decision, if it is correct, will immediately cause the majority of the clique to choose correctly. However, note that as clique size grows, the probability that the initial decision is incorrect also grows. It is therefore not obvious that the asymptotic fraction of the clique that chooses correctly approaches 1: If the initial decision is incorrect, the majority of the clique will be undecided after incorporating social information from the initial decision (See Fig. 8.1c). However, the social information exchange described above can still lead the remainder of the clique to overcome the impact of the wrong initial decision. Numerical simulations suggest that the the fraction of the clique that makes the correct choice does limit to 1, at least in the cases we have examined.

**Fig. 8.1.**
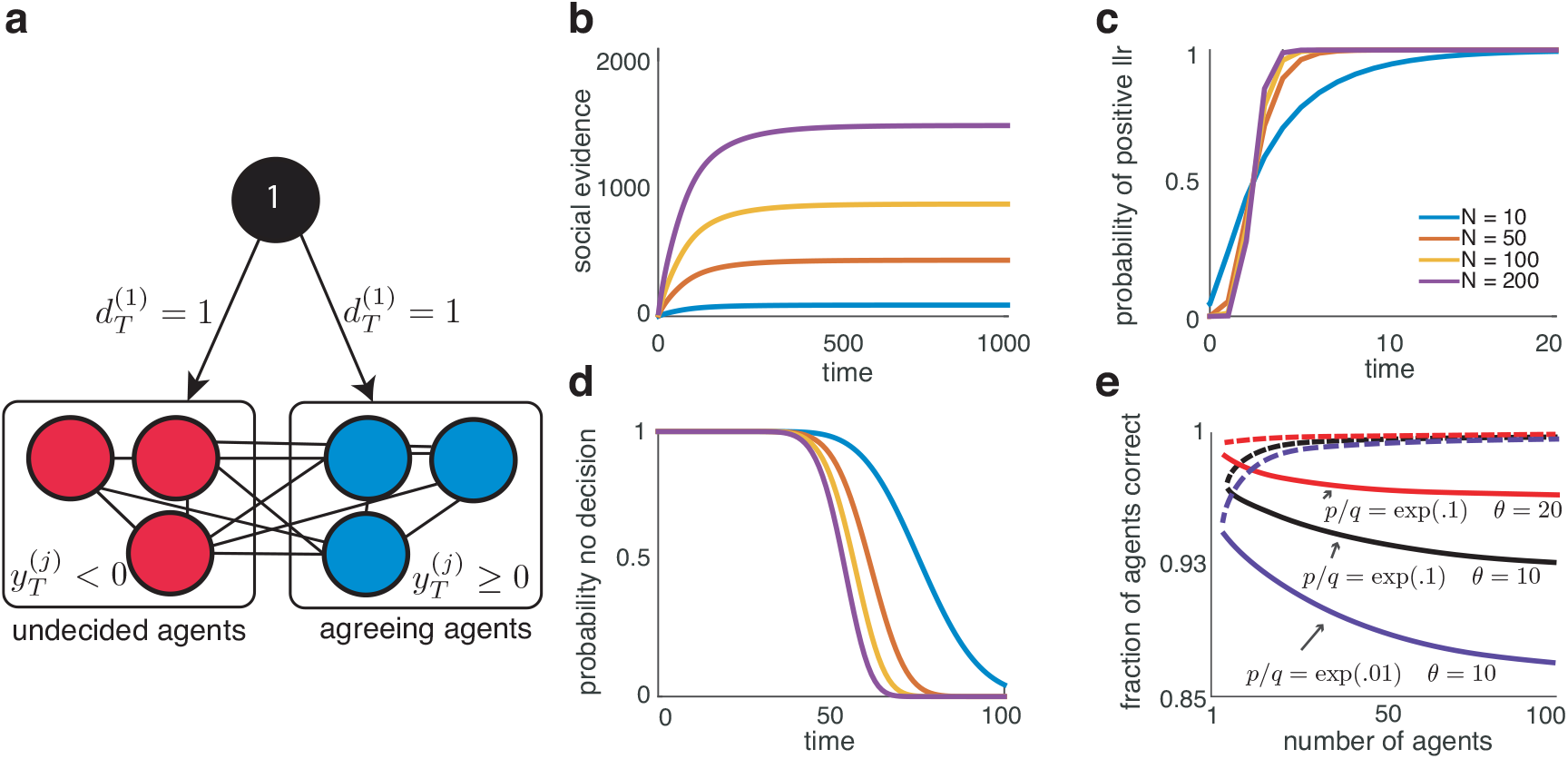
(a) In a clique of size N, we assume agent 1 makes a decision which is observed by all other agents. Agents whose beliefs are positive (agreeing agents), follow the decision of this initial agent. The remaining undecided agents continue to exchange social information. (b) Average social information, E[(a_1_ – (u_1_ – 1))R_+_], available to undecided agents after observing the first wave of agreeing agents as a function of the first decision time for different clique sizes. (c) The probability that the majority of the group has positive belief as a function of the first decision time for different clique sizes. This is equivalent to the probability that the majority of the clique consists of agreeing agents when the first decision occurs. (d) The probability that no agent has made a decision as a function of decision time for different clique sizes. Here, |θ_±_ |= 20, p = e/10, and q = 1/10. (e) The fraction of agents choosing correctly after the first wave following a decision (solid) and after equilibration (dashed) as a function of clique size for various values of p/q and θ.

## 9. Discussion

There has been extensive work on the mathematics of evidence accumulation by individual agents [7,9,46,56,59]. The resulting models have been very successful in explaining experimental data and the dynamics of decision making [24, 55]. In parallel, economists have developed mathematical models of networks of rational decision makers [25, 38]. However, in the work of economists static models are the norm, and the temporal dynamics of decisions are not studied explicitly. In contrast, temporal dynamics have been extensively studied in the psychology literature, but predominantly in the case of subjects making decisions based on private evidence [32] (although see [3, 26, 29]). Here, we provide a bridge between these research areas by developing a normative models of evidence accumulation in social networks.

The beliefs of evidence-accumulators in a network have been modeled by diffusively coupled drift-diffusion equations [44, 52]. Such models approximate continuous information-sharing between agents [52], and have lead to a number of interesting insights. However, it is not always clear when linear coupling between neighbors approximates the normative process of evidence accumulation and exchange between rational agents. In a related model, an agent observing a choice was assumed to experience a jump in its belief [10]. This study examined how the size of this jump affects the decisions of the collective. While not normative, this model displays interesting dynamics, and provides insight into decision making when agents over- or under-weight available evidence [6].

The belief dynamics of agents in our normative model differs from past accounts in several respects. First, we assume agents only communicate decision information, and do so normatively. In this case, coupling between agents is nonlinear and depends on a threshold-crossing process. We have shown that agents can exchange social evidence even in the absence of a decision when their measurement distributions or decision thresholds are not symmetric. Observing the decision state of a neighbor may result in further exchanges of social information with other neighbors. Furthermore, agents that do not have a direct view of all other agents in the network must marginalize over all possible decision states of the unobserved nodes. This can lead to complex computations, in contrast to the model discussed in [28], where a central agent sees all actions and optimizes a final decision assuming all other agents accumulate evidence independently.

The absence of choice is informative only in the case of asymmetries. This could be due to one choice being more rewarding than another [27, 39, 57]. For instance, the random dot motion task, which requires subjects to determine the dominant direction of randomly moving dots, has been extensively used in psychophysical and neurobiological studies. In this task subjects set their decision thresholds symmetrically when the reward and frequency of each choice is symmetric [24, 50]. However, when one drift direction is more likely or rewarded more, subjects exhibit accuracy and response time biases consistent with an asymmetric prior or thresholds [27, 39]. Recordings suggest neural activity representing decisions is dynamically modulated during the evidence accumulation process to reflect this bias [27, 43]. This is notably different from the static offsets apparent in the Bayesian model, but further investigation is needed to determine how priors are represented across the entire decision-making system [53].

In our model, unless threshold asymmetries are large, the social information communicated by the indecision of a single agent is weak. However, in large social networks the totality of such evidence provided by all other agents can be large compared to the private information obtained by an agent. Therefore, in large networks even small asymmetries can produce social information from indecisions that strongly influences decisions: Consider a new, desirable product that hits the market. If a large fraction of a social network is indecisive about purchasing the product, this can communicate potential issues with the product. This signal could be particularly strong if the product is particularly desirable upon its release. Another example is the case of decentralized detection: Consider a distributed network of independent environmental hazard sensors which only communicate their decision (e.g., whether the environment is hazardous or safe) to a central agent (fusion center), which combines their independent decisions [11]. The central agent only uses social information, but has a much more accurate estimate of the environmental state than any single sensor. Moreover, if decision thresholds are set asymmetrically, say so that the hazard threshold is much lower, the presence of indecisions in all sensors would suggest they are collecting evidence in favor of a safe environment. By computing the LLR of the survival probability from each independent sensor, the central agent could obtain strong evidence suggesting a safe environment even before receiving any decisions.

Our normative model is unlikely to accurately describe the decision making process of biological agents. The rationality of humans and other animals is bounded [51], while some of the computations that we have described are quite complex, and provide only marginal increases in the probability of making a correct choice. Thus biological agents are likely to perform only those computations – perhaps approximately – that provide the largest impact [13]. A normative model allows us to identify which computations are important, and which offer only fractional benefits.

A strong simplifying assumption in this work is that observers make uncorrelated measurements. This assumption will not hold in most situations, as members of a social group are likely to obtain dependent information. However, experimental evidence suggests that humans do not take into account such correlations when making decisions [17, 31] and behave as if the evidence gathered by different members of their social group is independent of their own. The types of models we describe may approximate the decision-making process of agents that make such assumptions and the impact of ignoring such correlations when they are present is an interesting avenue for future work. Our assumption that all agents in the network are identical can be relaxed. As long as all agents know each other’s measurement distributions and decision thresholds, social information can be computed as we have done in the present work, although notation becomes more cumbersome. The case in which agents do not know their neighbors’ measurement distributions and thresholds is considerably more complex. In this case agents either marginalize over the unknown parameters of their neighbors, or try to infer them based on the temporal dynamics of their decision states.

Our findings are distinct, but related to previous work on herding and common knowledge. In this modeling work agents typically make a single private measurement, followed by an exchange, or propagation of social information [5, 33]. In recurrent networks, agents can announce their preference for a choice, until they reach agreement with their neighbors [2, 22, 23, 36]. This framework can be simplified so that agents make binary decisions, based solely on their neighbors’ opinion, admitting asymptotic analyses of the cascade of decisions through networks with complex structure [60]. In contrast, we assume that agents accumulate private and social evidence, and make irrevocable decisions. The resulting decision-making processes are considerably different: For instance, in our model agents, and there is no guarantee that social learning occurs. Combining private and social evidence also makes it difficult to derive exact results, but we expect asymptotic formulas are possible for large cliques, and simpler assumptions on the decision process.

Key to understanding the collective behavior of social organisms is uncovering how they gather and exchange information to make decisions [13, 20]. Mathematical models allow us to quantify how different evidence-accumulation strategies impact experimentally observable characteristics of the decision-making process such as the speed and accuracy of choices [8], and the level of agreement among members of a collective [12]. Such models can thus lead the way towards understanding the decisions and disagreements that emerge in social groups.

## Acknowledgements

This work was supported by an NSF/NIH CRCNS grant (R01MH115557) and an NSF grant (DMS-1517629). BK and KJ were supported by NSF grant (DMS-1662305). KJ was also supported by NSF NeuroNex grant (DBI-1707400). ZPK was also supported by an NSF grant (DMS-1615737).

## Appendix A. Decomposition of an Agent’s Belief into Private and Social Information

The following proposition shows that the belief of each agent is a sum of information from private measurements, and observed decisions, as claimed in Eq. (3.2).

### Proposition A.1.

*Assume that in the network depicted in Fig. 3.1 agent 1 makes a choice at time T* ^(1)^*. If agent 2 has not yet made a decision, its belief is*

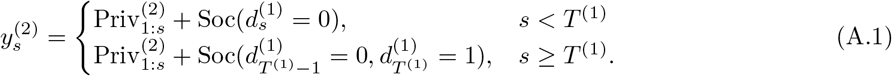

*Proof*. The decisions of agent 1 are independent from the observations of agent 2, when conditioned on the state, *H*. Thus

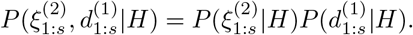

Hence, taking the logarithm of the ratio of 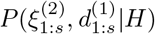 gives

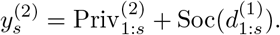

It remains to show that 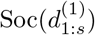 simplifies to the form given in Eq. (A.1).

Consider 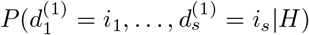 where *i*_1_ *, …, i*_s_ ∈ {*−*1, 0, 1}. Before agent 1 makes a decision, it communicates a decision state of 0, so if *s < T*^(1)^:

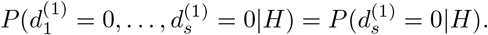

This equality holds because 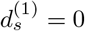 implies that all previous decision states must also be 0.

Similarly, when agent 1 makes a decision at time *T* ^(1)^, we can write

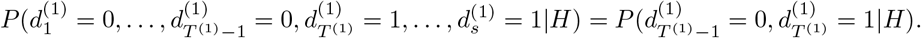

Again, this is because 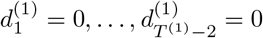 is implied by 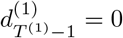 and the values of the decision states after time *T* ^(1)^, 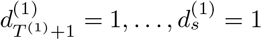 are implied by 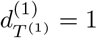. Thus

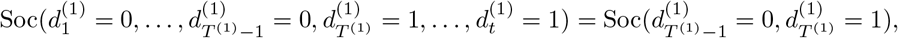

and we note that the evidence from the decision state depends on the value of the decision state and the time when it was first non-zero, i.e., when agent 1 made a choice.

## Appendix B. Proof of Proposition 4.1: Bounding Social Information from a Decision

We prove the case *d_T_* (1) = +1 in detail. The *d_T_* (1) = *−*1 is analogous. With Θ = (*θ_−_, θ*_+_), we have

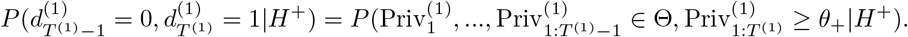

We observe that 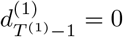 and 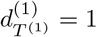 imply 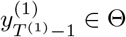 and 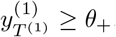. Hence

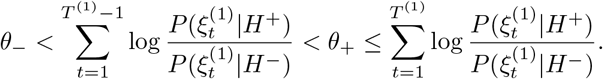

Following [7], the last of these inequalities implies

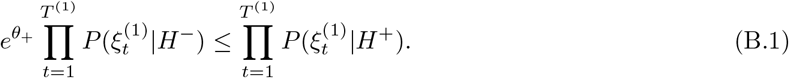

We define the set of *legal* chains of beliefs as those that lead to a decision at time *T* ^(1)^, but not earlier,

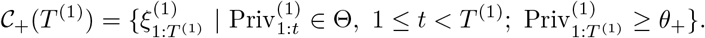

Note that *C*_+_(T^(1)^) ⊆ Ξ^T^(1)^^. Hence

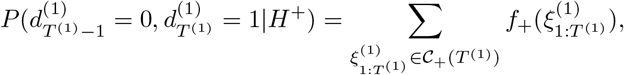

where we are using the notation

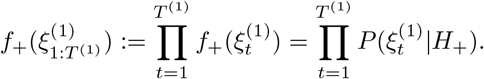

Every legal chain satisfies the inequality (B.1), so summing yields

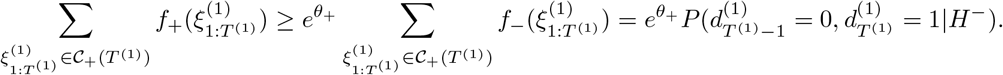

Thus

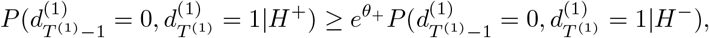

and so

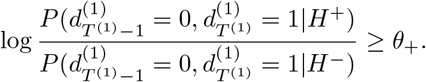

By noting 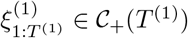, we can directly compute

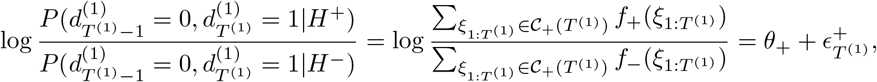

where we have defined

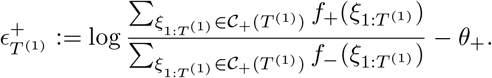

We can obtain a simpler upper bound by noting that for any 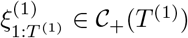, we have

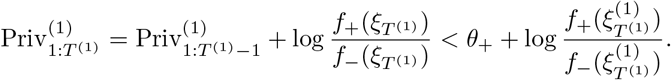

which implies

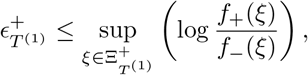

where we define

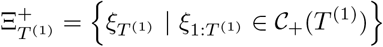

by positivity of log(*f*_+_(*ξ_T_*_(1)_)*/f_−_*(*ξ_T_*_(1)_) triggers a decision, and 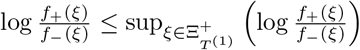 for all 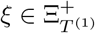 A similar argument can be used to derive bounds on 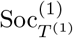 for 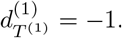.

## Appendix C. Explicit Formula for Social Information for Four-State Biased Random Walk

We use standard first-step analysis [54] to calculate non-decision social evidence for *θ*_+_ = 2 and *θ_−_* = *−*1. In this case we have four belief states, 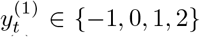, where 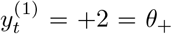 and 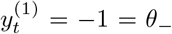 are absorbing boundaries. Let 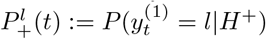. Then, for 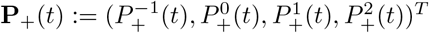,

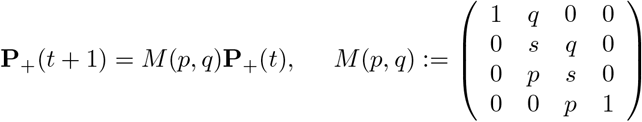

with initial condition 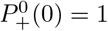 and *s* = 1 *− p − q*.

Let **v**(*t*) be a vector of probabilities that 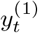 is at a non-absorbing state, then we can solve for

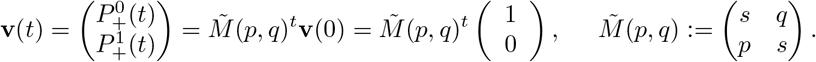

*M̃(p,q)* has eigenvalues 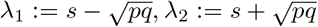 with eigenvectors

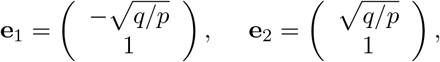
so that

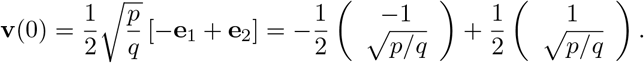

Thus,

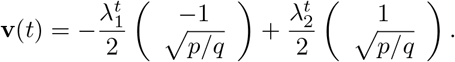

We calculate the eigenpairs for *M* ̃(*p, q*)^*t*^by swapping *p* and *q,* and can thus obtain an expression for **v**(*t*) in that case. Then,

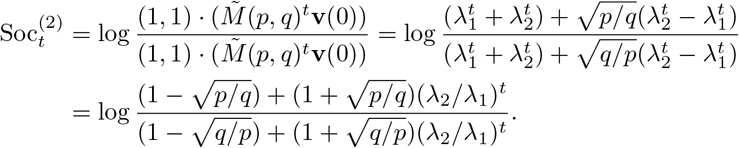

For 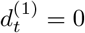 we have

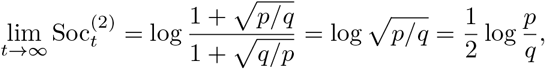

since |λ_2_| > λ_1._ For our choice of parameters, 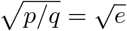 so

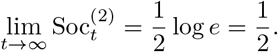

Note the recurring appearance of the quantity *p/q* in this calculation. This is a measure of the noisiness of the independent observations of the environment and is a proxy for the signal-to-noise ratio (SNR). In particular, note that if *p* ⪢ *q*, then 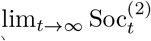 is large. On the other hand, if observations are extremely noisy, i.e. *p ≈ q*, then 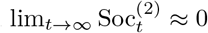

Interestingly, in the limit where all observations are informative (*s →* 0), the social information will alternate indefinitely between two values. In this case, *λ*_2_*/λ*_1_ *→ −*1, and we have

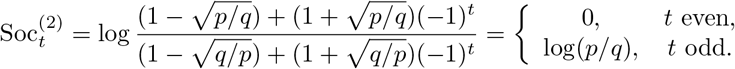

This follows from the fact that the belief of agent 1 must be at lattice site 0 (1) after an even (odd) number of observations.

## Appendix D. Proof of Proposition 5.2: Bounding Social Information from a Decision in a Recurrent Two-Agent Network

We use an argument similar to that in Proposition 4.1. First, consider the case in which *n* = 0, so private observation 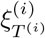 triggers the decision. If 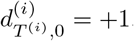, we know

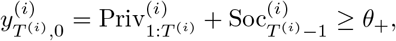

and since 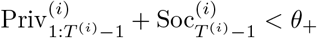, then

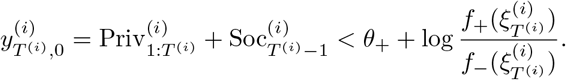

Marginalizing over all chains *C*_+_(*T* ^(*i*)^, 0) such that 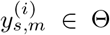 for all 0 *< s < T* ^(*i*)^, and corresponding 0 ≤ *m* ≤ *N*_*s*_ preceding the decision 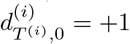 at (*T^(i)^, 0*), we thus find

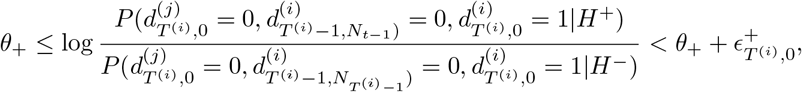

where *N_T_*^(*i*)^*−*1 is the maximal substep in the social information exchange following the private observation at timestep *T*^(*i*)^ *−* 1. A similar argument shows that

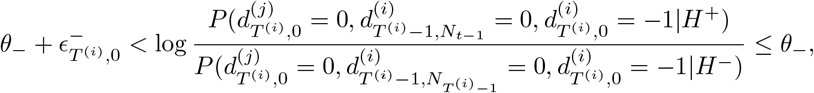

where

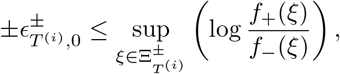

where we define

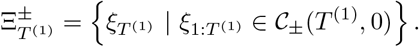

On the other hand, suppose 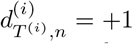, and 0 < *n* ≤ *N_T(i)_* so that the decision is reached during the social information exchange following an observation. Then,

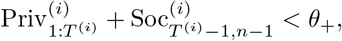

which implies

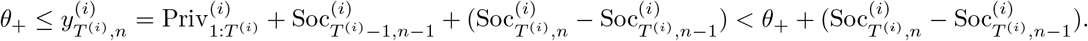

Marginalizing over all chains *C*_+_(*T* ^(*i*)^*, n*) such that 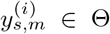 for all 0 *< s ≤ T* ^(*i*)^, and corresponding 0 *≤ m ≤ N_s_*, preceding the decision at (*T* ^(*i*)^*, n*),

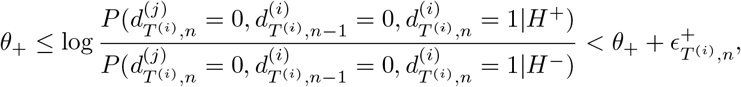

and similarly for *C_*(*T^(i)^, n*), we have

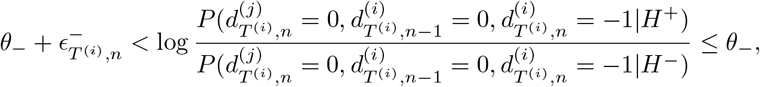

where

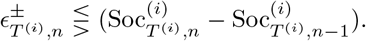

Following the arguments in Theorem 5.1, we note then that

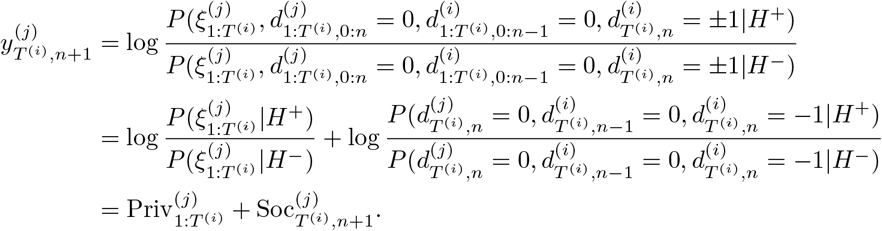

## Appendix E. Three agents on a directed line

Here we show that when agent 3 observes a decision by agent 2, at time *T* ^(2)^ the increment in 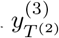 exceeds *θ,* depending mostly on the thresholds *θ,* but less on the informativeness of individual observations.

First, we note that for any *s = 1,* …, *T*^(2)^ − 1,

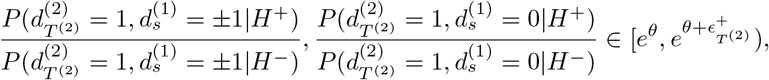

using arguments similar to those presented in Prop. 4.1 and Prop. 5.2. Therefore, the best estimate of these likelihood ratios can only bound them within finite intervals. However, such considerations muddle the presentation, and in fact in the case for which the measurement distributions *f*_+_(*ξ*) and *f_−_*(*ξ*) are close, max_*ξ*_|*f*_+_(*ξ*) *− f_−_*(*ξ*)| < *ϵ*, then 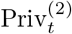 are small and so 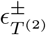 are also small. With this in mind, we write

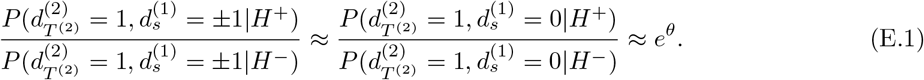

This is a common approximation in the literature on sequential probability ratio tests [58]. The situations where a decision from agent 1 does *not* immediately cause a decision from agent 2, conditioned on *H*^+^, are described by

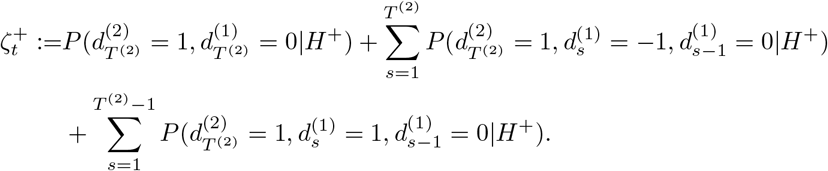

Note, when agent 2 makes an *H^+^* decision immediately after 1 makes an *H*^−^ decision, 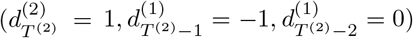, it must have done so due to private information 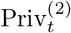 arriving immediately before the social information 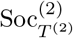 due to agent 1’s opposite decision, which would otherwise drive 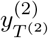 far from the +*θ* threshold. Since we have assumed 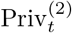 are small, this situation is extremely unlikely, so 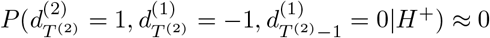. Using Eq. (E.1), we have

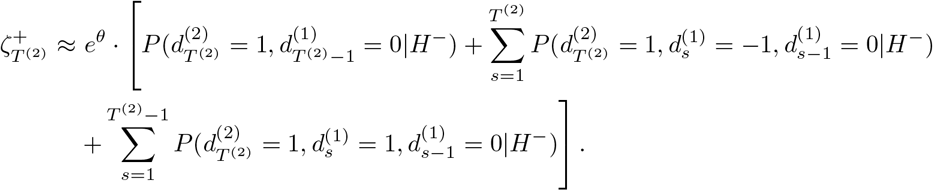

By Eq. (6.4), we have

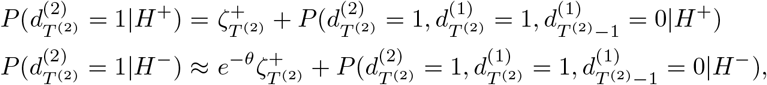

So that

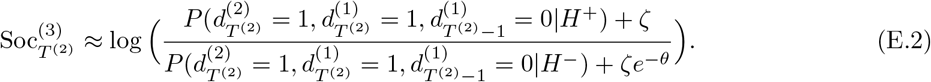

Note that

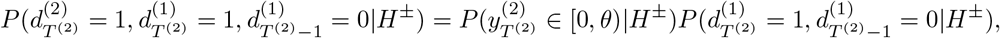

and

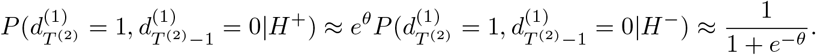

Let

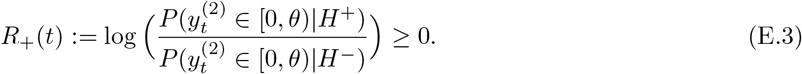

Note that we can increase *R*_+_(*t*) by increasing *θ*, without changing the measurements, or their distributions.

It follows that,

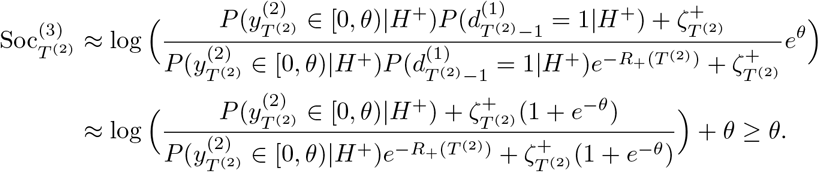

We therefore see that increasing *R*_+_(*T*^(2)^) increases the magnitude of social information received from observing a choice by agent 2.

The impact of a decision 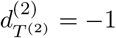 can be computed in a similarly, yielding

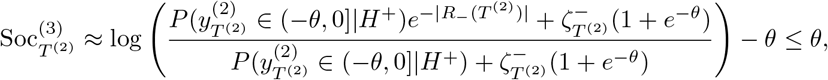

Where

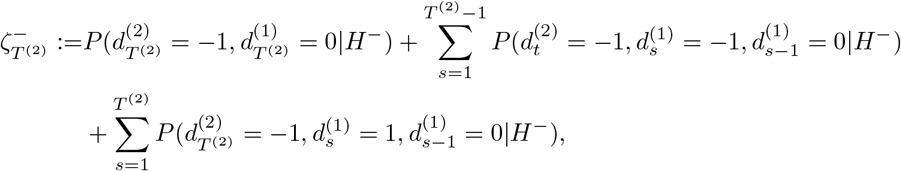

and

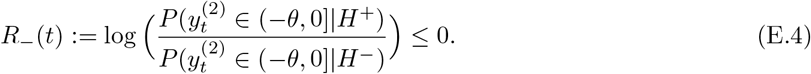

## Appendix F. Social Information from Bounds on Neighbor’s Beliefs

We next show that knowing that an agent’s belief lies within a subset of the interval Θ = (–*θ, θ*) provides at most an amount of social evidence equal to *θ*.

In the proof we assume that a sequence of private observations results in a belief (LLR) that can lie exactly on the decision threshold *θ*, as in the examples in the text. The proof can be extended to cases when information gained from individual measurements is small, so that 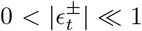. The proof in this case is equivalent, but somewhat more involved.

### PROPOSITION F.1.

*Let –θ < a* ≤ *b < θ. If agent j has not made a decision and has accumulated only private information by time T, then*

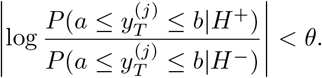

*Proof*. Define the subset *V_T_* (*a, b*) ⊂Ξ^*T*^of the product space of observations Ξ^*T*^consisting of observation sequences that result in a belief contained in [*a, b*] at time *T* :

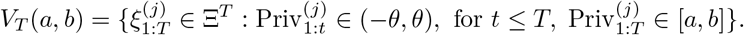

By definition, we can write

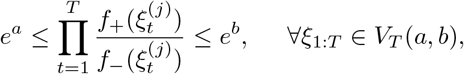

which can be rearranged as

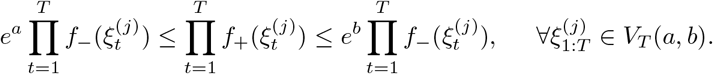

Summing over all 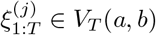 then yields

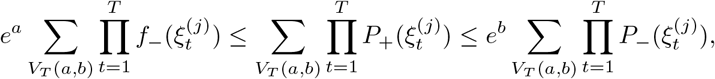

so that by noting

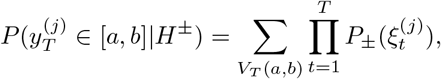

and rearranging we find

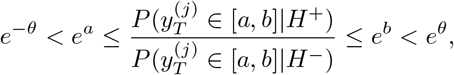

which implies the desired result. □

## Appendix G. Proving Reflection Symmetry of Social Information from LLR Bounds

### PROPOSITION G.1.

*Assume agent i has not received any social information at time t, so their belief, 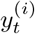, is based on private information. Also assume that the decision thresholds, θ_±_* = *±θ, and measurement distributions,f*_+_(*ξ*) = *f_−_*(*−ξ*)*, are symmetric. Let*

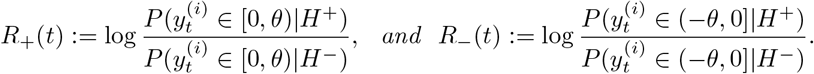

*Then R*–(*t*) = – *R*_+_(*t*).

*Proof*. Following from the argument in Proposition F.1, we can compute

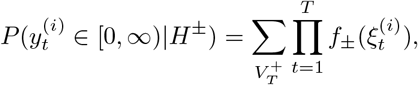

where we have defined

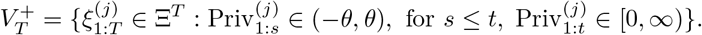

By symmetry, we know that for any 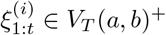, there exists 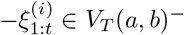 where

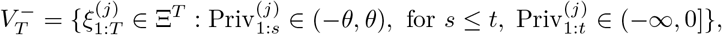

and vice versa. Therefore we can write

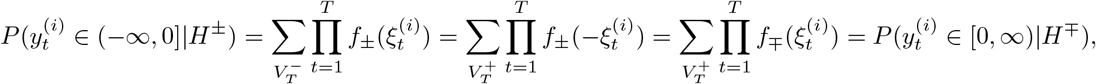

where the penultimate equality holds since *f*_+_(*ξ*) = *f_−_*(*−ξ*).

Therefore,

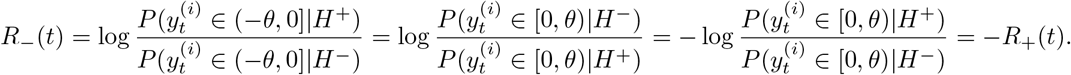

## Appendix H. Agreement Information in Cliques of Size

*N*. Let *A*_1_ denote the set of agreeing agents and *U*_1_ denote the set of undecided agents after the initial decision. We want to know what an undecided agent 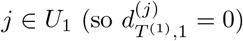 learns by observing the distribution of *A*_1_ and *U*_1_. To compute the evidence update for agent *j* first note that,

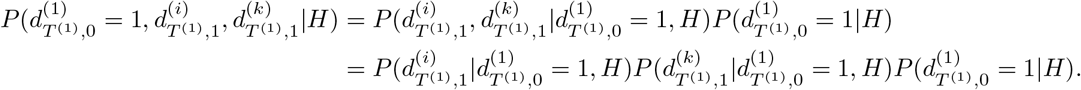

for any pair of agents *i ≠ j* different from 1. Therefore,

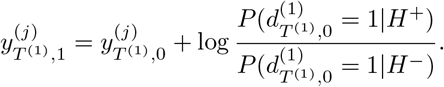

After observing this first wave of decision, all remaining undecided agents update their belief as,

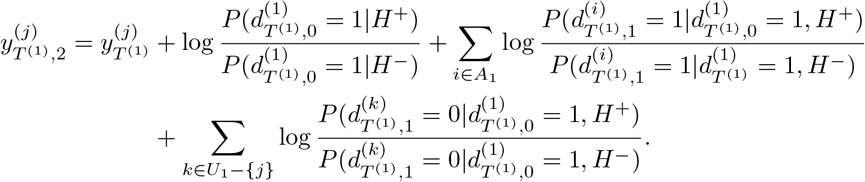

By conditional independence, this simplifies to:

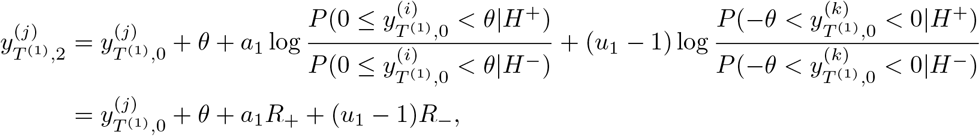

where *R_±_* are defined as in Eqs. 7.3 and E.3 . By symmetry, *R_−_*(*T* ^(1)^) = *−R*_+_(*T* ^(1)^). Thus

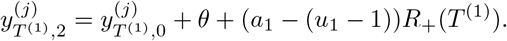

## Footnotes

1 In the remainder of this work we use the same measurement distributions in all examples. We always choose *p, q, s* so that decisions based solely on private evidence result in a LLR exactly at threshold.

